# Mutation in F-actin Polymerization Factor Suppresses Distal Arthrogryposis Type 5 (DA5) PIEZO2 Pathogenic Variant in *Caenorhabditis elegans*

**DOI:** 10.1101/2023.07.24.550416

**Authors:** Xiaofei Bai, Harold E. Smith, Luis O Romero, Briar Bell, Valeria Vásquez, Andy Golden

## Abstract

The mechanosensitive PIEZO channel family has been linked to over 26 disorders and diseases. Although progress has been made in understanding these channels at the structural and functional levels, the underlying mechanisms of PIEZO-associated diseases remain elusive. In this study, we engineered four PIEZO-based disease models using CRISPR/Cas9 gene editing. We performed an unbiased chemical mutagen-based genetic suppressor screen to identify putative suppressors of a conserved gain-of-function variant *pezo-1[R2405P]* that in human *PIEZO2* causes distal arthrogryposis type 5 (DA5; p. R2718P). Electrophysiological analyses indicate that *pezo-1(R2405P)* is a gain-of-function allele. Using genomic mapping and whole genome sequencing approaches, we identified a candidate suppressor allele in the *C. elegans* gene *gex-3.* This gene is an ortholog of human *NCKAP1*(NCK-associated protein 1), a subunit of the Wiskott-Aldrich syndrome protein (WASP)-verprolin homologous protein (WAVE/SCAR) complex, which regulates F-actin polymerization. Depletion of *gex-3* by RNAi, or with the suppressor allele *gex-3(av259[L353F])*, significantly restored the small brood size and low ovulation rate, as well as alleviated the crushed oocyte phenotype of the *pezo-1(R2405P)* mutant. Auxin-inducible degradation of GEX-3 revealed that only somatic-specific degradation of GEX-3 restored the reduced brood size in the *pezo-1(R2405P)* mutants. Additionally, actin organization and orientation were disrupted and distorted in the *pezo-1* mutants. Mutation of *gex-3(L353F)* partially alleviated these defects. The identification of *gex-3* as a suppressor of the pathogenic variant *pezo-1(R2405P)* suggests that the cytoskeleton plays an important role in regulating PIEZO channel activity and provides insight into the molecular mechanisms of DA5 and other PIEZO-associated diseases.

## Introduction

PIEZO proteins are ion channels that transduce mechanical stimuli into physiological responses. PIEZO1 and PIEZO2 trimers form a propellor-like structure with C-terminal domains of each PIEZO monomer assembling to create the central ion pore ^1, 2^. Dysfunction in both human PIEZO1 and PIEZO2 cause a variety of physiological disorders and diseases in humans ^3^. Gain-of-function mutations in PIEZO1 cause hereditary xerocytosis/dehydrated stomatocytosis (DHS) in red blood cells ^4–7^, while loss-of-function mutations in PIEZO1 could lead to congenital lymphatic dysplasia ^8^. In addition, pathologies in PIEZO2 gain-of-function and loss-of-function diseases mainly occurred in the distal limb and neuronal systems, which lead to joint contractures, muscle atrophy, and proprioception deficits ^9–11^. These pathogenic effects may occur due to insufficient or excessive responses of PIEZO channels to mechanical stimuli. At the cellular level, dysfunctional PIEZO channels may disrupt intracellular calcium homeostasis and interfere with downstream calcium-dependent signaling pathways ^3, 12^. Despite the electrophysiological characterization of PIEZO channel variants, at the functional level, the molecular and cellular mechanisms of PIEZO-associated diseases remain elusive.

The *C. elegans* spermatheca is a multicellular tube that can stretch and dilate. Oocytes enter the spermatheca through a distal neck, are fertilized within a central bag, and exit the spermatheca through the spermatheca-uterine (sp-ut) valve. The spermathecal valves and bag cells are spatiotemporally coordinated to control oocyte entry during ovulation, as well as oocyte exit after fertilization in the spermatheca. The contractility of the spermathecal tissue is driven by the coordinated activation of myosin ^13–15^. After ovulation, the fertilized oocytes are expelled from the spermatheca and into the uterus, pushing many sperm with it. These sperm must navigate back into the spermatheca for subsequent ovulations. The oocytes and somatic sheath cells secrete chemoattractant molecules and prostaglandins into the uterus to guide the sperm back to the spermatheca ^16, 17^. The *pezo-1* gene is the sole ortholog of human *PIEZO1* and *PIEZO2* in *C. elegans*. Dysfunctional PEZO-1 resulted in reduced brood size and low ovulation rate, as well as sperm guidance and navigational defects ^18^. Previous studies have shown that cytoskeletal components play a crucial role in coordinating the contractibility of the spermathecal cells^13–15^. Furthermore, PIEZO channels activity has been found to be regulated by the cytoskeleton integrity^34–35, 37, 40, 60–61^. Based on these findings, we hypothesize that the PIEZO channel plays a functional role during ovulation and contractility within the spermathecal tissue by regulating cytoskeletal activity, specifically actin polymerization. However, the exact role of the PEZO-1 channels in governing the cytoskeletal activity during ovulation remains elusive.

In this study, we established a disease model of human PIEZO channels by generating orthologous variants in the *C. elegans pezo-1* gene using CRISPR/Cas9 gene editing. All homozygous *pezo-1* pathogenic variants reduced brood size and ovulation rate. Two of the mutants, including *pezo-1(R2405L)* and *pezo-1(R2405P)*, caused sperm attraction defects, in which a portion of the sperm failed to migrate back to the spermatheca. Using a forward genetic screen, we found a suppressor allele in the gene *gex-3* which belongs to the *NCKAP1*(NCK associated protein 1), a subunit of Wiskott-Aldrich syndrome protein (WASP)-verprolin homologous protein (WAVE/SCAR) complex. Depletion of *gex-3* by RNAi and the suppressor allele *gex-3(L353F)* alleviated the small brood size and low ovulation rate, as well as the crushed oocyte phenotypes of the *pezo-1(R2405P)* mutant. Using an auxin-inducible degradation (AID) system, we drove tissue-specific degradation of GEX-3 in the *pezo-1(R2405P)* background. Reduced brood size was only restored in the somatic-specific degradation strain, suggesting that the loss-of-*gex-3* in somatic tissues primarily contributed to the suppression of *pezo-1* gain-of-function mutation *(R2405P)*. Lastly, we found that actin organization was disrupted and distorted in the *pezo-1* mutants, and that *gex-3(L353F)* partially alleviated actin defects in the *pezo-1(R2405P)* background. Overall, we used *C. elegans* to study the pathophysiological effect of an ortholog human *PIEZO1* mutation causing DA5 (*pezo-1 R2405P)* and identified the putative suppressor *gex-3(L353F)*. Here, we provide new insights into the nature of PIEZO channels and their interaction with cytoskeletal organizational elements.

## Results

### PIEZO pathogenic variants cause reproductive deficiency in *C. elegans*

To study underlying molecular mechanisms of PIEZO-associated diseases, we used CRISPR/Cas9 gene editing to generate four disease-relevant PIEZO alleles in the C-terminal domains of *C. elegans pezo-1*, the sole worm ortholog of human *PIEZO1* and *PIEZO2* (Fig. 1A). Each of the alleles were orthologous mutations associated with three different diseases: dehydrated hereditary stomatocytosis (DHS), distal arthrogryposis subtype 5 disorder (DA5), and Gordon syndrome (GS) (Table 1). All homozygous *pezo-1* mutations caused a 27-81% reduction in brood size compared to the wild type control (Fig. 1B). Particularly, non-conservative amino acid residue mutations *pezo-1(R2405P)* and *pezo-1(R2405L)* decreased brood size by 70%, while a more conservative (polar) mutation *pezo-1(R2405Q)* decreased the brood size by ∼20% (Fig. 1B).

**Figure 1.**
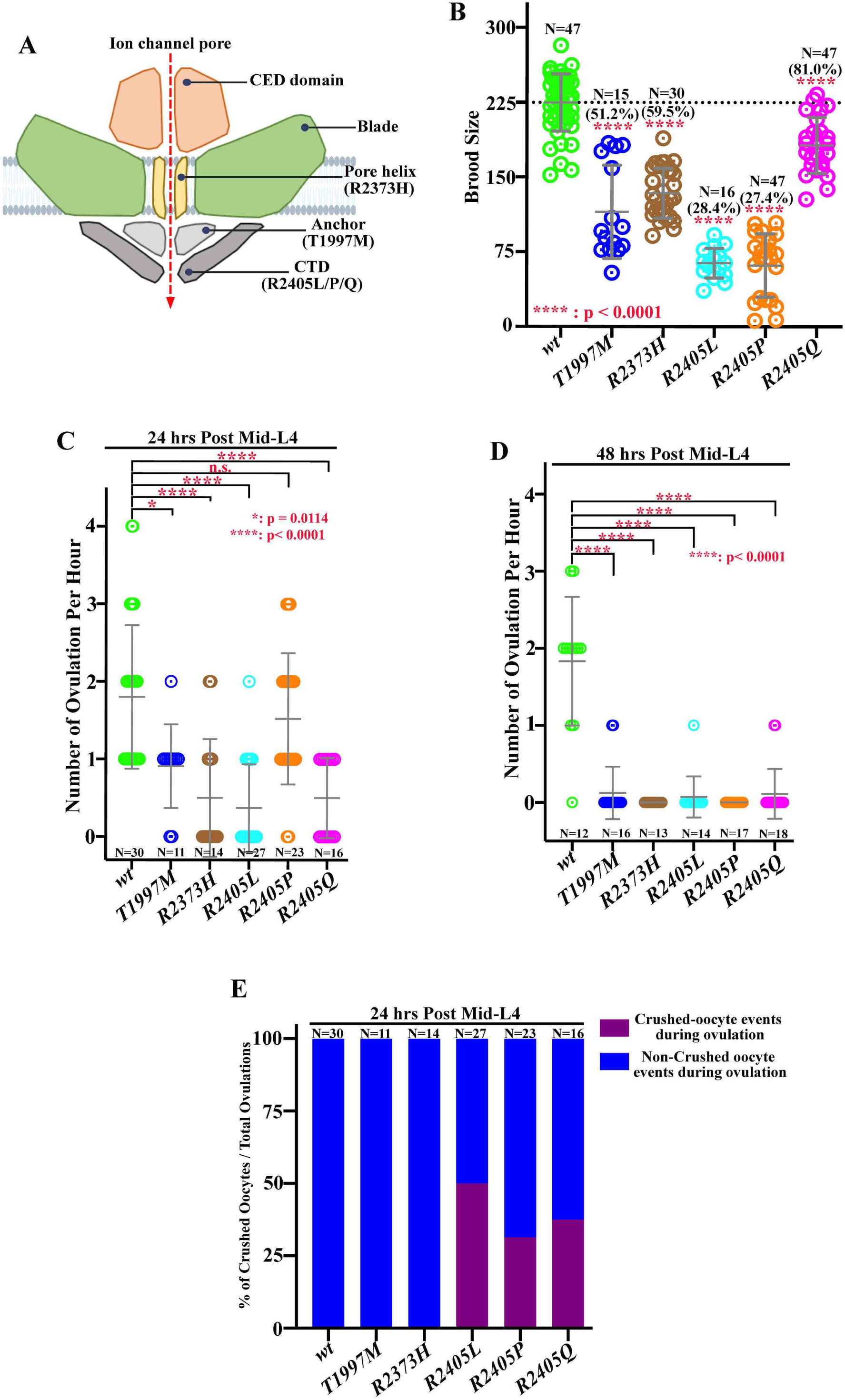
PIEZO pathogenic variants caused reproductive deficiency in *C. elegans*. (A) Diagram of the location of the PEZO-1 pathogenic residues in the PEZO-1 channel used in this study. (B) Brood size was reduced in all tested *pezo-1* pathogenic mutants when compared with wildtype. % in () indicate % of wildtype brood size. N values indicated the tested animals in (B). (C-E) Quantification of the oocyte ovulation rate and percentage of crushed oocytes of wildtype and *pezo-1* mutants during ovulation at different ages. N values indicated the number of the tested gonad in (C-E). The oocyte ovulation rate was significantly reduced in the *pezo-1* mutant adults. One-way ANOVA test (B-D) or Chi-squared-test (E). P-values: *: p= 0.0114; ****, p<0.0001 (t-test).

**Table 1.** Summary of domains and diseases caused by each pathogenic variant.

| **PEZO-1** | **hPIEZO1** | **Channel Domains** | **Disease** | **hPIEZO2** | **Channel Domains** | **Disease** |
| --- | --- | --- | --- | --- | --- | --- |
| T1997M | T2127M | Anchor | DHS<sup>6</sup> | T2356M | Anchor | DA5<sup>54</sup> |
| R2373H | R2456H | Pore helix | DHS<sup>54</sup> | R2686H | Pore helix | GS/DA5<sup>54</sup> |
| R2405L | R2488L | CTD | N/A | R2718L | CTD | DA5<sup>55</sup> |
| R2405P | R2488P | CTD | N/A | R2718P | CTD | DA5<sup>55</sup> |
| R2405Q | R2488Q | CTD | DHS<sup>7</sup> | R2718Q | CTD | N/A |

Dehydrated hereditary stomatocytosis (DHS), Distal Arthrogryposis type 5 (DA5), and Gordon syndrome (GS). [#] Reference Number.

Our previous study demonstrated that the reduced brood size in *pezo-1* mutants was due to lower ovulation rate as well as oocyte crushing while exiting the spermathecal-uterine (sp-ut) valve ^18^. To test whether the ovulation was affected in each of the current *pezo-1* mutants, we performed live imaging to record the ovulation process for at least one hour and analyzed the ovulation performance in each mutant. Indeed, we determined a low ovulation rate in all tested *pezo-1* mutants (Fig. 1C-D). Of note, the ovulation rate was reduced to nearly zero in older animals (day-2 adults), while wild-type controls continued to ovulate ∼1.83 times per hour per gonad arm (Fig. 1D). We also observed that approximately 30% of the ovulated oocytes from day 1 *pezo-1(R2405L/P/Q)* variants were crushed when transiting through the sp-ut valve, while no crushed oocytes were observed in the wild-type animals (Fig. 1E). Overall, our data suggest that *pezo-1* pathogenic variants caused a reduction in brood size, low ovulation rates, and crushed oocytes.

### PIEZO pathogenic variants disrupt sperm guidance in *C. elegans*

Various *pezo-1* null mutants exhibited defective sperm guidance ^18^. Either self-sperm or male sperm failed to navigate back to the spermatheca after each ovulation, thus depleting the spermathecal sperm. To test whether each *pezo-1* missense mutant fails to attract sperm back to the spermatheca, male sperm navigational performance was assessed *in vivo* by staining wild-type male sperm with a vital fluorescent dye, MitoTracker CMXRos, which efficiently stains sperm mitochondria in live animals ^17^. The stained wild-type males were placed for 30 minutes together with the label-free *pezo-1* mutant hermaphrodites. We isolated the hermaphrodites and allowed the sperm to navigate through the uterus to the spermatheca for one hour. We quantified the sperm distribution by counting the number of fluorescent-labeled sperm in each zone (Fig. 2A-G) ^18^. In wild-type hermaphrodites, 90.3% of fluorescent sperm navigated through the uterus and accumulated in the spermatheca (Fig. 2B-B’, H). The sperm distribution rate in the *pezo-1(T1997M)*, *pezo-1(R2373H)*, and *pezo-1(R2405Q)* hermaphrodites were identical to wild-type worms [93.3 % for *pezo-1(T1997M)*, 98.2% for *pezo-1(R2373H)*, 79.4% for *pezo-1(R2405Q)*] (Fig. C-D’, G-G’ and H). However, in the *pezo-1(R2405L)* and *pezo-1(R2405P)* reproductive tracts, the fluorescence-labeled sperm displayed defective sperm navigation. Only 56.1% and 57.5% of the labeled sperm in the *pezo-1(R2405L)* and *pezo-1(R2405P)* hermaphrodites were able to migrate back to zone 3, which was the zone adjacent to, and including the spermatheca (Fig. 2 E-F’, H), while the rest of the labeled sperm failed to reach the spermatheca and remained throughout zones 1 and 2, which are furthest from the spermatheca (Fig. 2 E-F’, H).

**Figure 2.**
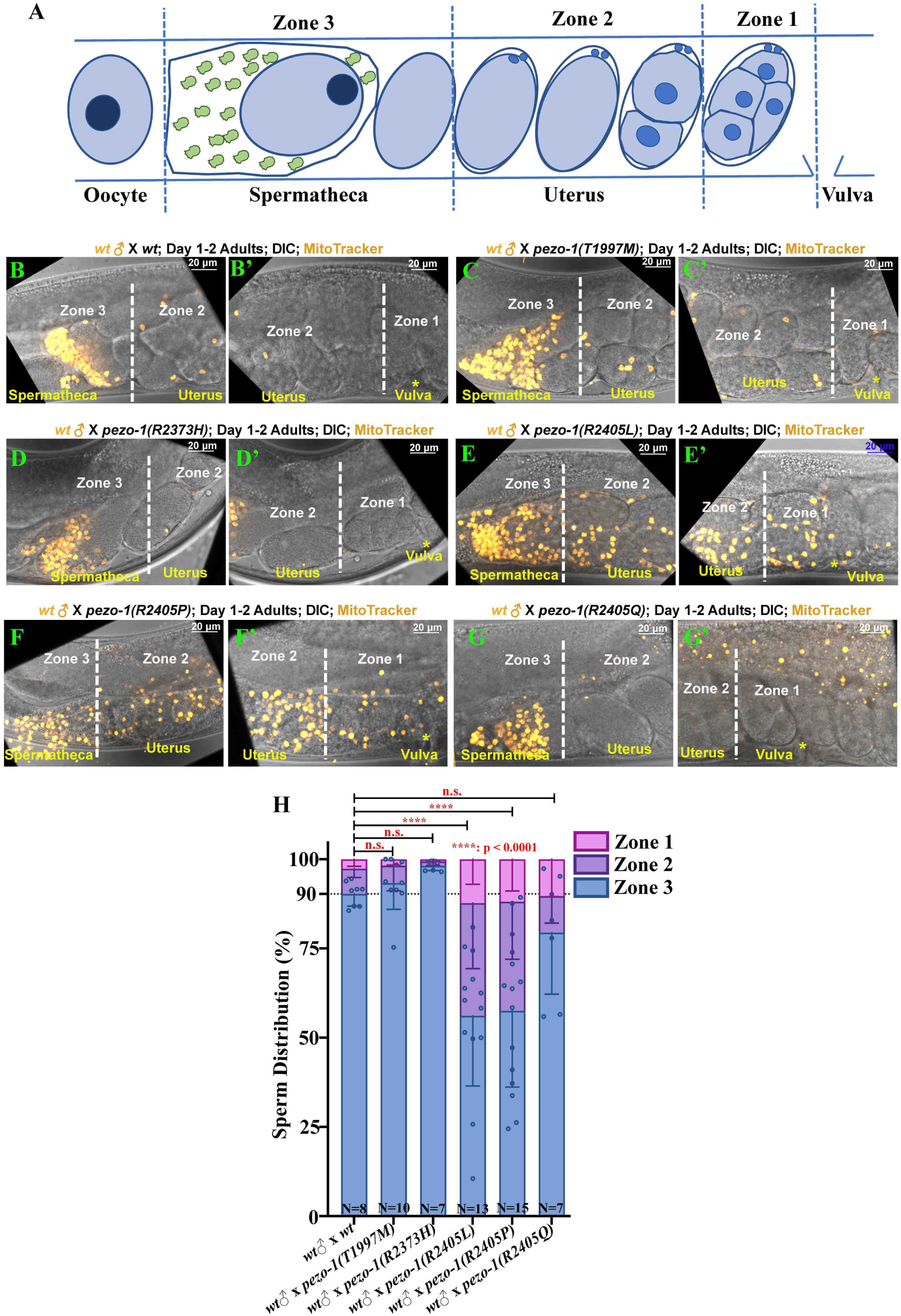
Sperm guidance and navigation is disrupted in PIEZO pathogenic variants. (A) To quantify sperm migration, sperm distribution was counted in three zones, including Zone 3 (A), which is the spermatheca region and the space containing the +1 fertilized embryo, whereas Zone 1 (A) is the area closest to the vulva, and Zone 2 (A) is the area between Zone 1 and Zone 3. Sperm distribution is measured 1 hr after the Mitotracker labeled males were removed from the mating plate. (B-G’) The distribution of fluorescent male sperm (yellow dots) labeled with MitoTracker in the three zones in both wildtype and *pezo-1* mutants. Yellow asterisks indicate the vulva (B’, C’, D’, E’, F’, and G’). Scale bars are indicated at top right in each panel. (H) Quantification of sperm distribution values of wild type and each *pezo-1* mutant. One-way ANOVA test (H). N numbers indicate the tested animals. P-values: ****, p<0.0001 (t-test).

### *pezo-1(R2405P)* is a gain-of-function allele

The *pezo-1(R2405P)* allele caused the most severe defects in the *C. elegans* reproductive tract (Fig. 1-2). This mutation is equivalent to the mouse PIEZO1 R2514 mutation, which resides at the end of the α1 helix of the C-terminal domain. We have previously determined that the longest isoform of *pezo-1* (isoform G; wormbase.org v. WS280) encodes a mechanosensitive ion channel ^19^. Of note, the behavior observed in the *pezo-1(R2405P)* mutant was similar to various pezo-1 null mutants previously described 18, suggesting that loss- or gain-of-function (LOF and GOF, respectively) *pezo-1* mutations are equally pathogenic, similar to their human counterparts 9-11. To determine the effect of *pezo-1(R2405P)* on PEZO-1 channel function, we heterologously expressed *C. elegans* wild-type and *pezo-1(R2405P)* mutant constructs in a *Spodoptera frugiperda* (Sf9) cell line and measured their mechanical response in the whole-cell patch-clamp configuration while stimulating with a piezo-electrically driven glass probe (Fig. 3A-C). Current densities for the *pezo-1(R2405P)* mutant were larger than non-infected cells, although not significantly different than wild-type *pezo-1* (Fig. 3D). Noteworthy, substituting an arginine with a proline at position 2405 results in mechanocurrents that inactivate more slowly than the wild-type channel, as reflected by the large time constant of inactivation (Fig. 3E). We also determined that the R2405P construct requires less mechanical stimulation to open than wild-type *pezo-1* (Fig. 3F). Taken together, our results suggest that *pezo-1(R2405P)* is a gain-of-function allele as PEZO-1 takes longer to stop conducting cations inside the cells.

**Figure 3.**
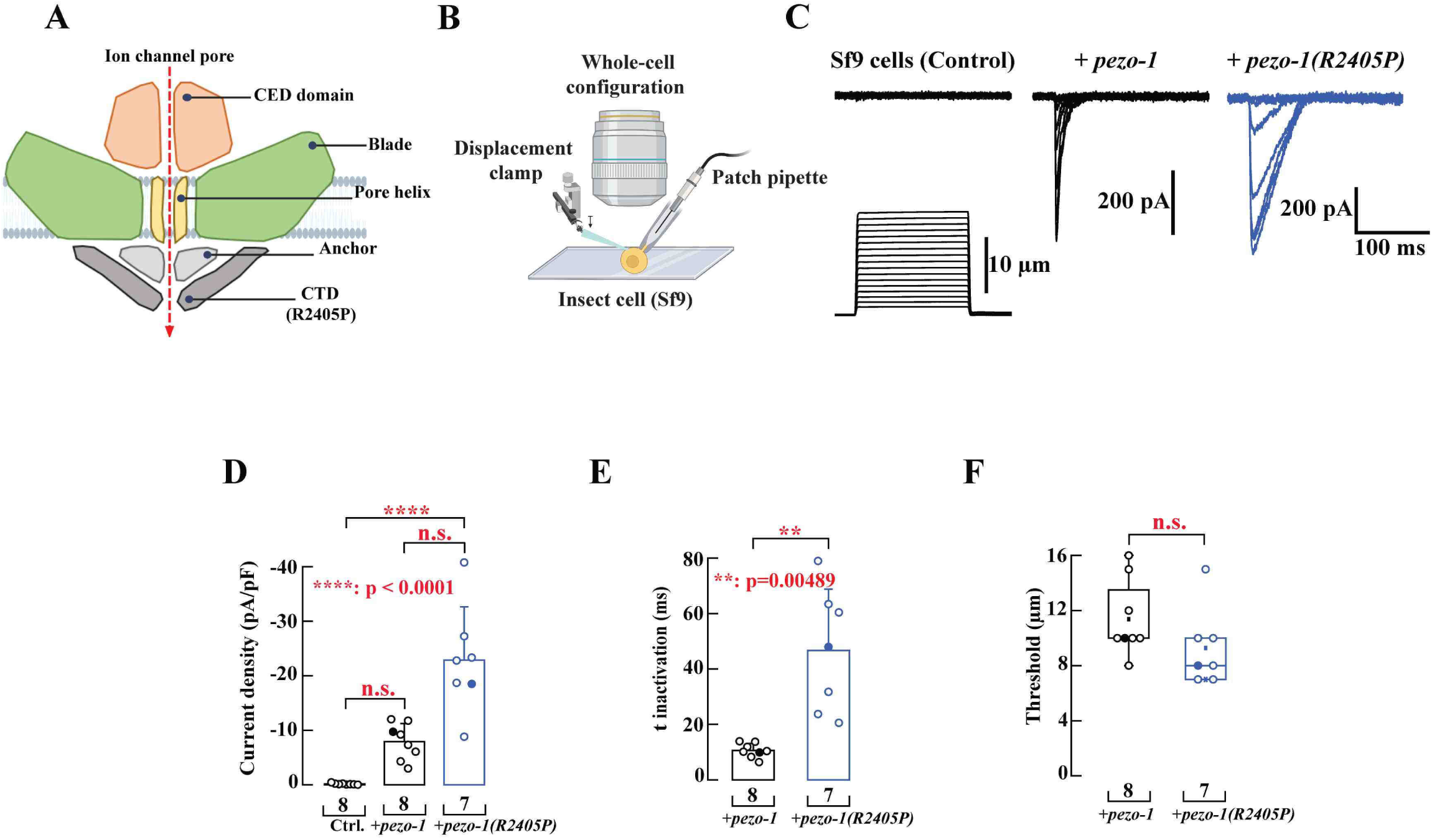
Electrophysiological characterization of PEZO-1 gain-of function mutation *pezo-1(R2450P)* (A) Diagram of the location of the PEZO-1(R2405P) pathogenic residue in the PEZO-1 channel. (B) Schematic representation of the mechanical stimulation poked by a blunt pipette applied to Sf9 cells infected with baculovirus containing *pezo-1* (WT or R2405P constructs) recorded in the whole-cell configuration. Created with BioRender.com. (C) Representative whole-cell patch clamp recordings (at −60 mV) of currents elicited by mechanical stimulation of Sf9 cells, uninfected (control), expressing *pezo-1* WT, or *pezo-1(R2405P)*. Sf9 cells were poked by a heat-polished blunt glass pipette (3–4 µm) driven by a piezo servo controller. Displacement measurements were obtained with a square-pulse protocol consisting of 1-µm incremental indentation steps. Recordings with leak currents >200 pA and with access resistance >10 MΩ, as well as cells with giga-seals that did not withstand at least five consecutive steps of mechanical stimulation, were excluded from analyses. (D) Current density elicited by maximum displacement (−60 mV) of Sf9 cells expressing *pezo-1* WT or *pezo-1(R2405P)*. Bars are mean ± SD. Kruskal-Wallis (H = 18.35; p = 0.0001) and Dunn’s multiple comparisons test. (E)Time constants of inactivation elicited by maximum displacement (−60 mV) of Sf9 cells expressing *pezo-1* WT or *pezo-1(R2405P)*. Bars are mean ± SD. Two-tailed unpaired t test with Welch correction (t = -4.29). (F) Boxplots show the displacement thresholds required to elicit mechanocurrents of Sf9 cells expressing *pezo-1* WT or *pezo-1(R2405P)*. Boxplots show mean (square), median (bisecting line), bounds of box (75^th^ to 25^th^ percentiles), outlier range with 1.5 coefficient (whiskers), and minimum and maximum data points. Two-tailed Mann-Whitney test (U = 13.5). Filled circles come from the representative traces shown in (C). n values were indicated under each column. Post-hoc p-values are denoted in the corresponding panels. P-values: **: p=0.0049; ****, p<0.0001 (t-test).

### EMS-based forward genetic screen to identify suppressors of *pezo-1(R2405P)*

The small brood size in the *pezo-1* mutants was a simple readout to perform a forward genetic screen to isolate genetic suppressors of PIEZO disease mutations. To better understand the mechanism by which PEZO-1 regulates nematode reproduction, we conducted a genetic suppressor screen of the gain-of-function mutant *pezo-1(R2405P),* which has the lowest brood size of all the alleles we tested. Previously, we showed that *pezo-1* interacted genetically with an ER calcium regulator SERCA pump (*sca-1*)^18^. To increase the sensitivity of our forward screening, we shifted the synchronized EMS-treated F2 population of *pezo-1(R2405P)* embryos to *sca-1* RNAi food and maintained the animals at 25°C until the candidate suppressor lines were isolated (Fig. S1A). Under these sensitive screening conditions, wild-type hermaphrodites produced viable animals with a significant reduction in brood size (Fig. S1B), while *sca-1* RNAi treatment of our *pezo-1(R2405P)* mutant led to sterility or extremely low brood sizes (greater than 80% reduction compared to wild-type worms) after 1-2 generations (Fig. S1B). We screened ∼150,000 haploid genomes and isolated one stable suppressor line which partially restored the reduced brood size (n=161.1) when compared with *pezo-1(R2405P)* or *pezo-1(R2405P)* on *sca-1* RNAi food for one generation (n= 87.4 and n= 34.6, respectively) (n= 34.6) (Fig. S1B).

### MIP-MAP mapping and suppressor allele validation

Using a high-throughput genomic mapping strategy, which involves the molecular inversion probes genomic mapping (MIP-MAP) strategy^20^, we were able to identify the genetic modifiers in the suppressor line *sup1* (Fig. S2). To maintain the *pezo-1(R2405P)* allele during the mapping process, we generated *pezo-1(R2405P)* in the mapping strain VC20019 background by CRISPR/Cas9 gene editing and named the strain *pezo-1(R2405P^MM^)* (Fig. S2A) ^20^. This strain appeared to have similar phenotypes to that of our original *pezo-1(R2405P)* mutant. We carefully pooled suppressed F2 progeny from the cross and allowed the F2 population to expand for ten generations for MIP-MAP analysis (Fig. S2A). Since the flanking regions of the modifiers originated from a *pezo-1(R2405P)* background, the occurrence frequency of the MIP-MAP probes at the loci responsible for suppression would drop to nearly zero ^20^. Theoretically, reading the frequency of MIP-MAP probes would allow us to narrow down the target regions bearing the putative. After MIP-MAP analysis and whole genome sequencing, we mapped a single genomic region in the suppressor line *pezo-1(R2405P); sup1* (Fig. S2B), which contained candidate genes located on Chromosome IV (9.1 -12.0 Mb) (Fig. S2).

### A mutation in the WAVE regulatory complex NCKAP1 suppresses the reproductive defects caused by *pezo-1(R2405P)*

Assuming the candidate suppressors are loss-of-function alleles, we used RNAi to deplete expression of these candidate genes in the *pezo-1(R2405P)* mutant. From an RNAi trial experiment, we identified a candidate modifier gene, *gex-3,* on Chromosome IV (Fig. 4A and Fig. S2B). *gex-3* is the ortholog of human *NCKAP1* (NCK-associated protein 1), one of the five WAVE regulatory complex subunits, which regulates the formation of the actin cytoskeleton via the Arp2/3 complex ^21^. To assess the knock-down efficiency of *gex-3* RNAi, we first fed *gex-3* (RNAi) bacteria to mNG::GEX-3 animals and quantified the fluorescent intensity of mNG::GEX-3 after 24 hours of RNAi treatment (Fig. S3) ^22^. The fluorescent signals of mNG::GEX-3 in the germline and spermatheca were dramatically reduced when compared with RNAi negative control (Fig. S3A-C), suggesting that *gex-3* RNAi was sufficient and specific to delete *gex-3* gene expression *in vivo*. We then depleted *gex-3* by RNAi in wild-type worms and all *pezo-1* pathogenic mutants (Fig. 4A-B) to validate its suppression. *gex-3*(RNAi) caused significant embryonic lethality in all tested strains (Fig. 4B); however, most embryos did hatch. *gex-3*(RNAi) also led to reduced brood sizes in both wild-type worms and the *pezo-1(T1997M)* mutant (Fig. 4A). The brood size in other tested mutants, including *pezo-1(R2373H)*, *pezo-1(R2405L)*, and *pezo-1(R2373Q)*, were not significantly affected by *gex-3*(RNAi). Of note, the reduced brood size was significantly restored in *pezo-1(R2405P)* (n= 113.1) by *gex-3*(RNAi) (n=179.3) (Fig. 4A). Additionally, depletion of *gex-3* by RNAi caused reduced embryonic lethality in the *pezo-1(R2405P)* mutant when compared with *gex-3* (RNAi) in the wild-type worms, indicating that *pezo-1(R2405P)* moderately suppressed the phenotype caused by *gex-3* (RNAi) (Fig. 4B). Furthermore, live imaging indicated that treatment with *gex-3* (RNAi) resulted in similar ovulation rates but fewer number of crushed oocytes on the *pezo-1(R2405P)* background (Fig. 4C-E). Using RNAi, we next tested whether other WAVE regulatory complex subunits suppressed the defects in the *pezo-1(R2405P)* mutant by depleting *abi-1* and *gex-1* (*wve-1*), orthologs of human Abelson interactor gene (*ABI1*) and human Wiskott-Aldrich Syndrome Protein Family Member 1 gene (*WASF1*) respectively, using RNAi. However, the smaller brood size of the *pezo-1(R2405P)* mutant was not restored with either RNAi treatment (Fig. S4A-B).

**Figure 4.**
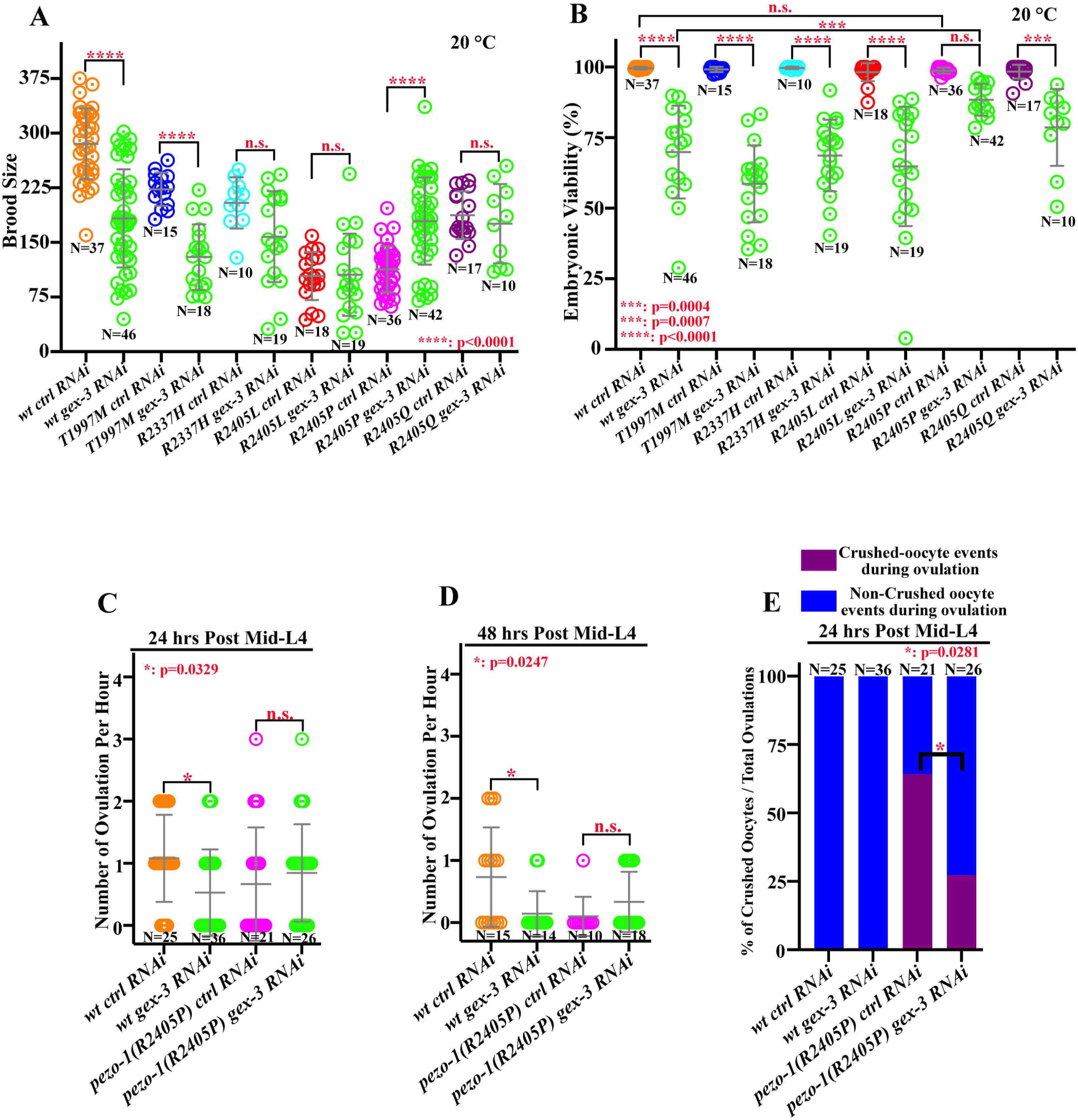
The WAVE regulatory complex NCKAP1 suppresses the reproductive defects in the *pezo-1(R2405P)* mutant. (A) *gex-3*(RNAi) treatment reduced brood size in wildtype and *pezo-1(T1997M)* mutant animals, while significantly restoring the brood size in *pezo-1(R2405P)* mutant. (B) depletion of *gex-3* by RNAi led to various levels of embryonic lethality in all tested animals, however, the *pezo-1(R2405P)* allele partially alleviated the lethality when compared to wildtype control. N values indicated the number of the tested animals in (A-B). (C-E) Quantification of the oocyte ovulation rate and percentage of crushed oocytes of wildtype and *pezo-1* mutants without or without *gex-3*(RNAi) treatment. N values indicated the number of the tested gonad in (C-E). One-way ANOVA test (A-D) or Chi-squared-test (E). P-values: *: p=0.0329 (C); *: p=0.0247 (D), *: p= 0.0281 (E); ***, p=0.0004 (B); ***, p=0.0007 (B); ****, p<0.0001 (A-B).

### mNG::GEX-3 expresses in multiple tissues and co-localizes with mScarlet::PEZO-1

We used a CRISPR knock-in green fluorescent reporter mNeonGreen::GEX-3 (mNG::GEX-3) to determine the subcellular localization of GEX-3 in *C. elegans* ^22^. mNG::GEX-3 was widely expressed from embryonic stages to adulthood (Fig. 5). Of note, the mNG::GEX-3 was strongly expressed in various tissues subjected to mechanical stimulation, including the pharyngeal-intestinal valve and spermatheca (Fig. 5A-B’’’). Notably, mNG::GEX-3 also co-colocalized with mScarlet::PEZO-1 at the pharyngeal-intestinal valve (Fig. 5A-A’’’), spermathecal membrane (Fig. 5B-B’’’), and early embryonic membranes (Fig. 5 C-C’’’).

**Figure 5.**
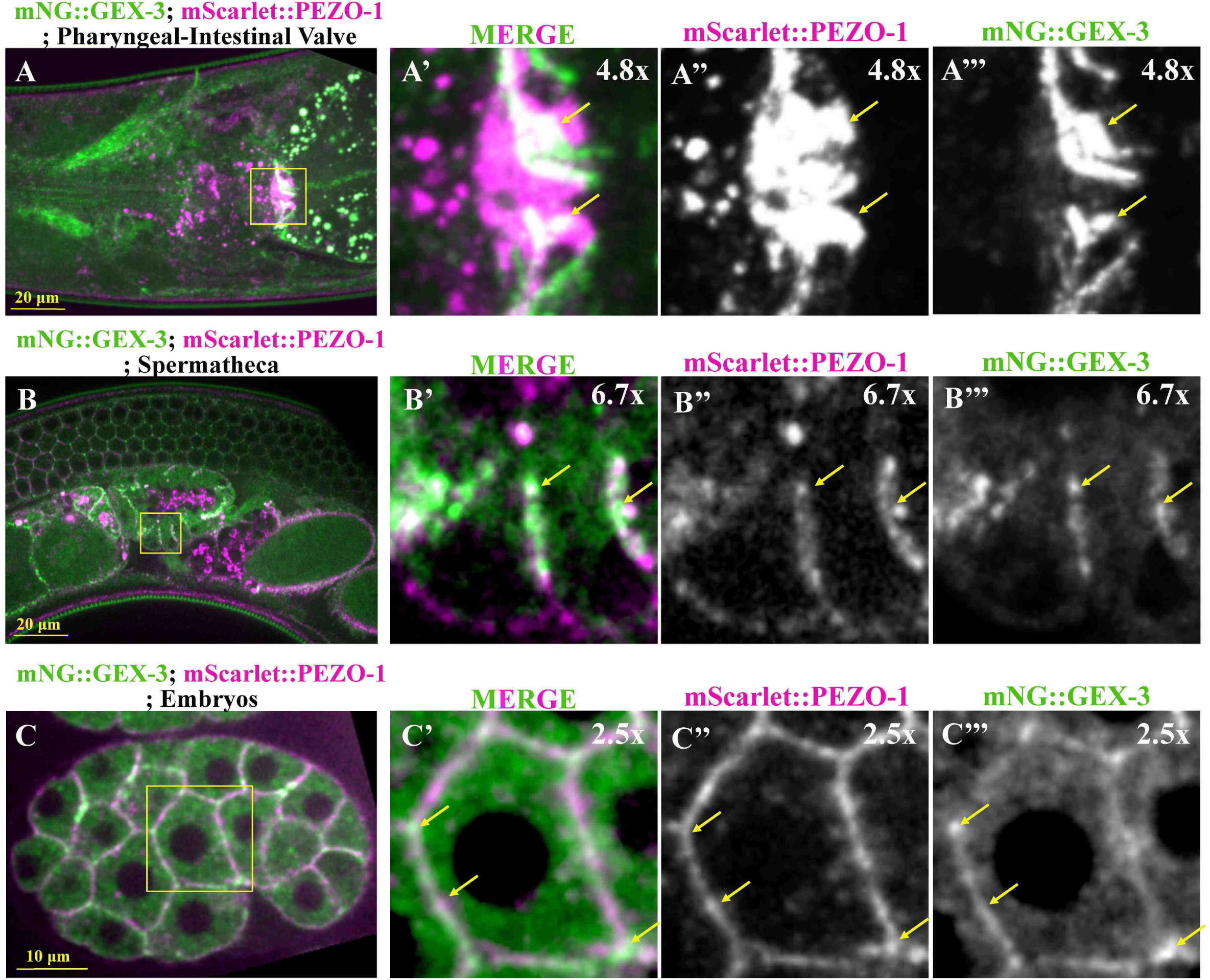
mScarlet::PEZO-1 colocalizes with mNG::GEX-3 in multiple tissues and cells. (A-A’’’) mScarlet::PEZO-1 (magenta in A-A’) colocalizes with mNG::GEX-3 (green in A-A’) at the pharyngeal-intestinal valve. The enlarged pictures of the yellow rectangle area indicated the colocalization of mScarlet::PEZO-1 (magenta in A’) and mNG::GEX-3 (green in A’). (B-B’’’) Colocalization of mScarlet::PEZO-1 (magenta in B-B’) and mNG::GEX-3 (green in B-B’) on the spermathecal membrane (yellow box in B-B’). Enlarged picture of yellow box in B showed colocalization of mScarlet::PEZO (magenta in B’) and mNG::GEX-3 (green in B’) on the membrane (yellow arrows). (C-C’’’) mNG::GEX-3 and mScarlet::PEZO-1 was observed at the plasma membrane of the early embryos. Both mNG::GEX-3 (green in C’) and mScarlet::PEZO-1 (magenta in C’) were expressed on the embryonic plasma membrane (yellow arrows). Colocalization of mNG::GEX-3 and mScarlet::PEZO-1 appears white color. Scale bar was labeled in each panel.

### *gex-3(L353F)* suppresses the reproductive defects caused by *pezo-1(R2405P)*

After confirming that the of loss-of-function of *gex-3* suppressed the phenotypes in *pezo-1(R2405P)* animals, we generated a candidate suppressor allele by changing Leu353 to Phenylalanine by CRISPR/Cas9 gene editing on the wild-type background. This allele was renamed as *gex-3(L353F)*. To assess its suppression during reproduction, this newly generated allele was then introduced onto the *pezo-1(R2405P)* background by CRISPR/Cas9 gene editing. The predicted GEX-3 structure from AlphaFold suggests that the residue Leu353 was located at the N-terminus of a helical segment (Fig. 6A) ^23, 24^. The homozygous *gex-3(L353F)* strain exhibited a smaller brood size when compared to wild-type worms (Fig. 6B). Unlike *gex-3(RNAi)* treatment, there was no embryonic lethality observed in the *gex-3(L353F)* mutant generated by CRISPR/Cas9 (Fig. 6C). To determine whether this missense mutation disrupted the temporal and/or spatial expression pattern of GEX-3, we generated the *gex-3(L353F)* allele using CRISPR/Cas9 on the mNG::GEX-3 reporter strain, which was named mNG::*gex-3(L353F)*. Endogenously tagged mNG::GEX-3 was expressed in multiple *C. elegans* tissues, including embryos, the germline, and pharynx (Fig. 5, S5A, and C). Similar localization patterns were found in the mNG::*gex-3(L353F)* mutant (Fig. S5B and D). Overall, these data suggest that the *gex-3(L353F)* allele display a weak effect, which may only partially compromise GEX-3 protein function, without altering trafficking and cellular localization of GEX-3 *in vivo*.

**Figure 6.**
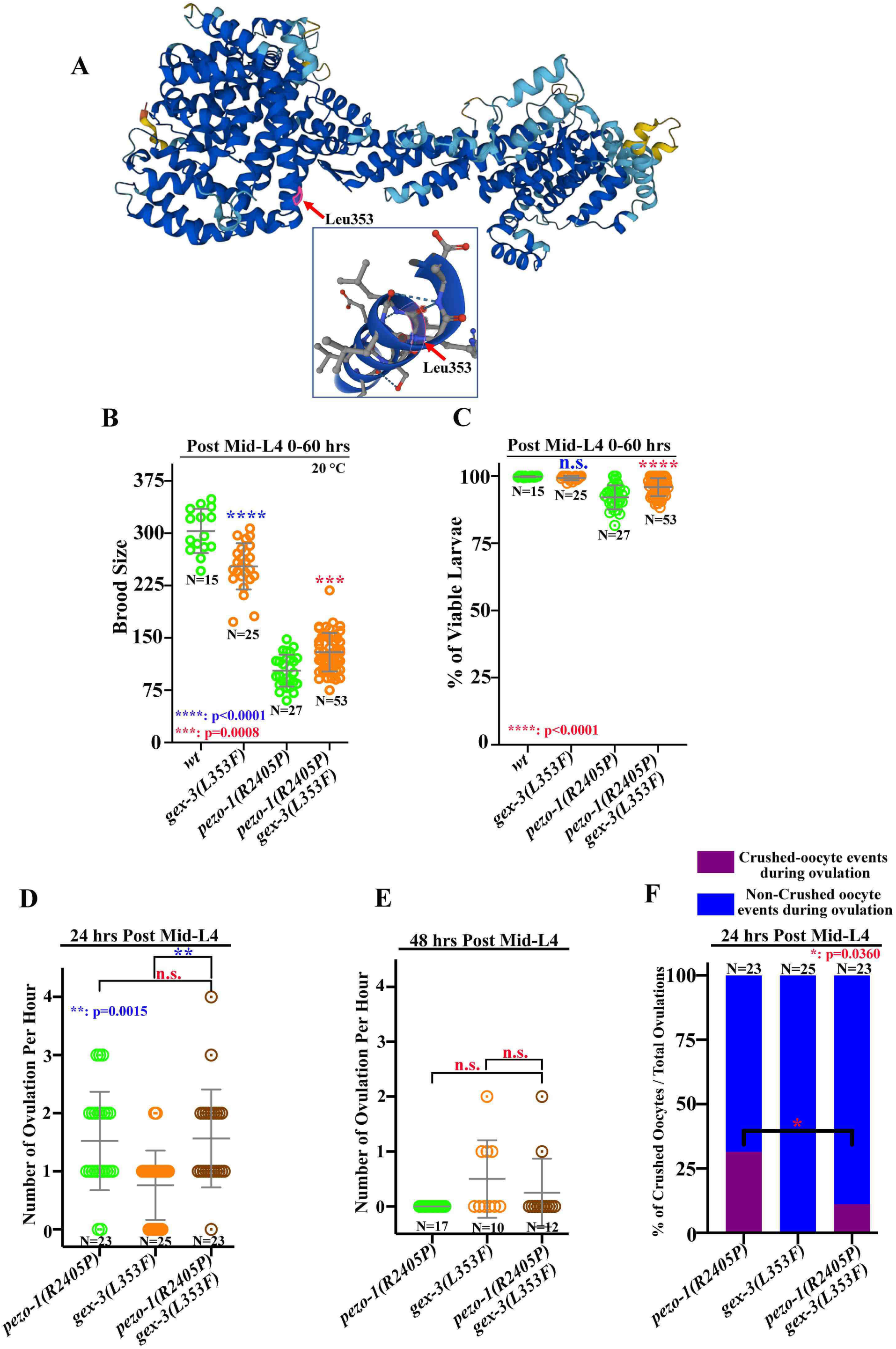
*gex-3(L353F)* suppresses the reproductive defects in the *pezo-1(R2405P)* mutant. (A) The structure of GEX-3 from Alphafold indicated the Leu353 residue was on the N-terminus of a helix (highlighted by red arrow). Different colors represent model confidence: dark blue Very high (pLDDT > 90); cyan-Confident (90 > pLDDT > 70); yellow-Low (70 > pLDDT > 50); orange-Very low (pLDDT < 50). (B) *gex-3*(L353F) reduced brood size in wildtype animals but suppressed the smaller brood size in the *pezo-1(R2405P)* mutant animals. (C) The embryonic lethality in *pezo-1(R2405P)*, *gex-3(L353F)* and double mutants. N values indicated the number of the tested animals in (B-C). (D-F) Quantification of the oocyte ovulation rate and percentage of crushed oocytes of wildtype, *pezo-1(R2405P)*, *gex-3(L353F)*, and double mutants at different ages. N values indicated the number of the tested gonad in (D-F). One-way ANOVA test (B-E) or Chi-squared-test (F). P-values: *, p=0.0360 (F); **, p=0.0015 (D); ***, p=0.0008 (B); ****, p<0.0001 (B-C).

The *gex-3(L353F); pezo-1(R2405P)* double mutant restored the reduced brood size and subtle embryonic lethality observed in the *pezo-1(R2405P)* mutant alone (Fig. 6B-C). In addition, the double mutant reduced the crushed oocyte rate (from 31.4% to 11.1%) and slightly increased the ovulation rate to (0.25+/-0.62) in the day-2 adults, as compared to *pezo-1(R2405P)* alone (0.0+/-0.0) (Fig. 6D-F). The double mutant displayed a higher ovulation rate in day-1 animals (1.57+/-0.84) than *gex-3(L353F)* alone (0.76+/-0.60) (Fig. 6D), suggesting that *pezo-1(R2405P)* and *gex-3(L353F)* mutually suppresses one another’s reproductive defects.

The small brood size of the *pezo-1(R2405P)* mutant might raise from the high number of sperm that fail to navigate back to the spermatheca. To test whether the *gex-3(L353F)* allele restored sperm attraction back to the spermatheca, we assessed the MitoTracker-stained male sperm navigational performance in both *gex-3(L353F)* and *gex-3(L353F); pezo-1(R2405P)*. In *gex-3(L353F)* hermaphrodites mated with stained wild-type males, over 90% of fluorescent sperm navigated through the uterus and accumulated in the spermatheca (Fig. S6A-A’, C) within one hour of mating. The sperm distribution rate in the double mutant hermaphrodites was 43.9%, which was similar to the *pezo-1(R2405P)* single mutant (57.5%, Fig. 2G), suggesting that *gex-3(L353F)* had no effect on sperm attraction behavior. Overall, our genetic interaction data suggests that both *gex-3* (RNAi) or the *gex-3(L353F)* mutant could partially suppress the reproductive defects of the *pezo-1(R2405P)* strain, but only at the ovulation and brood size levels. Of note, the depletion of *gex-3* by RNAi resulted in allele-specific suppression of the *pezo-1(R2405P)* mutant, but did not suppress other missense alleles tested. Meanwhile, the *pezo-1(R2405P)* allele also partially alleviated the embryonic lethality (Fig. 4B) and increased low ovulation rate (Fig. 4C) caused by *gex-3* (RNAi), suggesting that *pezo-1(R2405P)* and *gex-3* (RNAi) are mutual suppressors.

### Somatic tissue-specific degradation of GEX-3 suppresses the small brood size of the *pezo-1(R2405P)* mutant

GEX-3 is expressed in all reproductive tissues, including the spermatheca, germline, oocytes, and embryos (Fig. 5B-C). To better understand the role of GEX-3 in suppressing the subfertility of the *pezo-1(R2405P)* mutant, we used an auxin-inducible degradation system (AID) to degrade GEX-3 in somatic tissues or the germ line ^25^. We knocked-in a cassette with degron and GFP coding sequence at the *gex-3* N-terminus using CRISPR/Cas9 gene editing (named AID::GFP::GEX-3) so that the degron strain could be visualized and quantified by a GFP fluorescent signal (Fig. S7). The somatic and germline specific AID strains were driven by *Peft-3* and *Pmex-5,* respectively ^26^ and activated when the animals were exposed to 2 mM auxin (indole-3-acetic acid, IAA). The GFP fluorescence intensities in the induced animals were significantly reduced compared to vehicle control (Fig. S7B-B’’, D-D’’, and E-F). The strain expressing the degron interactor transgene *Pmex-5::tir-1::BFP::AID* led to a 2-3 fold reduction in fluorescence intensity of AID::GFP::GEX-3 in the germline and oocytes (Fig. S7B, B’’, and E), however, the intensity was not affected in the somatic tissues (Fig. S7E). The *Peft-3::tir-1::BFP::AID* led to approximate 1.5 fold reduction of fluorescent intensity of AID::GFP::GEX-3 in the sheath, spermathecal cells, and germline (Fig.S7D, D’’, and F), suggesting the somatic degron strain affected the AID::GFP::GEX-3 in both somatic and germline cells.

To assess the tissue-specific suppression of GEX-3 in *pezo-1(R2405P)*, we introduced the *pezo-1(R2405P)* allele into each degron strain by CRISPR/Cas9 gene editing. We exposed L4 animals to either 0.25% ethanol as a control or 1-2 mM IAA, then brood sizes and embryonic lethality rate were determined 0-60 hours post L4 (Fig. 7A-B). Interestingly, the brood size of the *pezo-1(R2405P)* mutant was significantly restored in the somatic tissue-specific AID::GFP::GEX-3 strain (Fig. 7A). Meanwhile, there were no significant changes in brood size in germline-specific AID::GFP::GEX-3 driven by the *Pmex-5* promoter (Fig. 7A). Additionally, depletion of GEX-3 in both somatic and germline tissues led to severe embryonic lethality (Fig. 7B). Unlike the partial rescue of embryonic lethality of *gex-3 (RNAi)* by *pezo-1(R2405P),* embryonic lethality was close to 100% for *pezo-1(R2405P)*; AID::GFP::GEX-3 strains, regardless of the promoter used for the *tir-1::BFP::AID* cassette (Fig. 7B). Therefore, degradation of GEX-3 in the somatic tissue suppressed the small brood size of *pezo-1(R2405P)*, likely due to the role of *gex-3* in the spermatheca or sheath contraction. This seems likely as we observed an improvement in ovulation performance when combining *gex-3* RNAi and *gex-3(L353F)* with *pezo-1(R2405P)* (Fig. 4 and 6).

**Figure 7.**
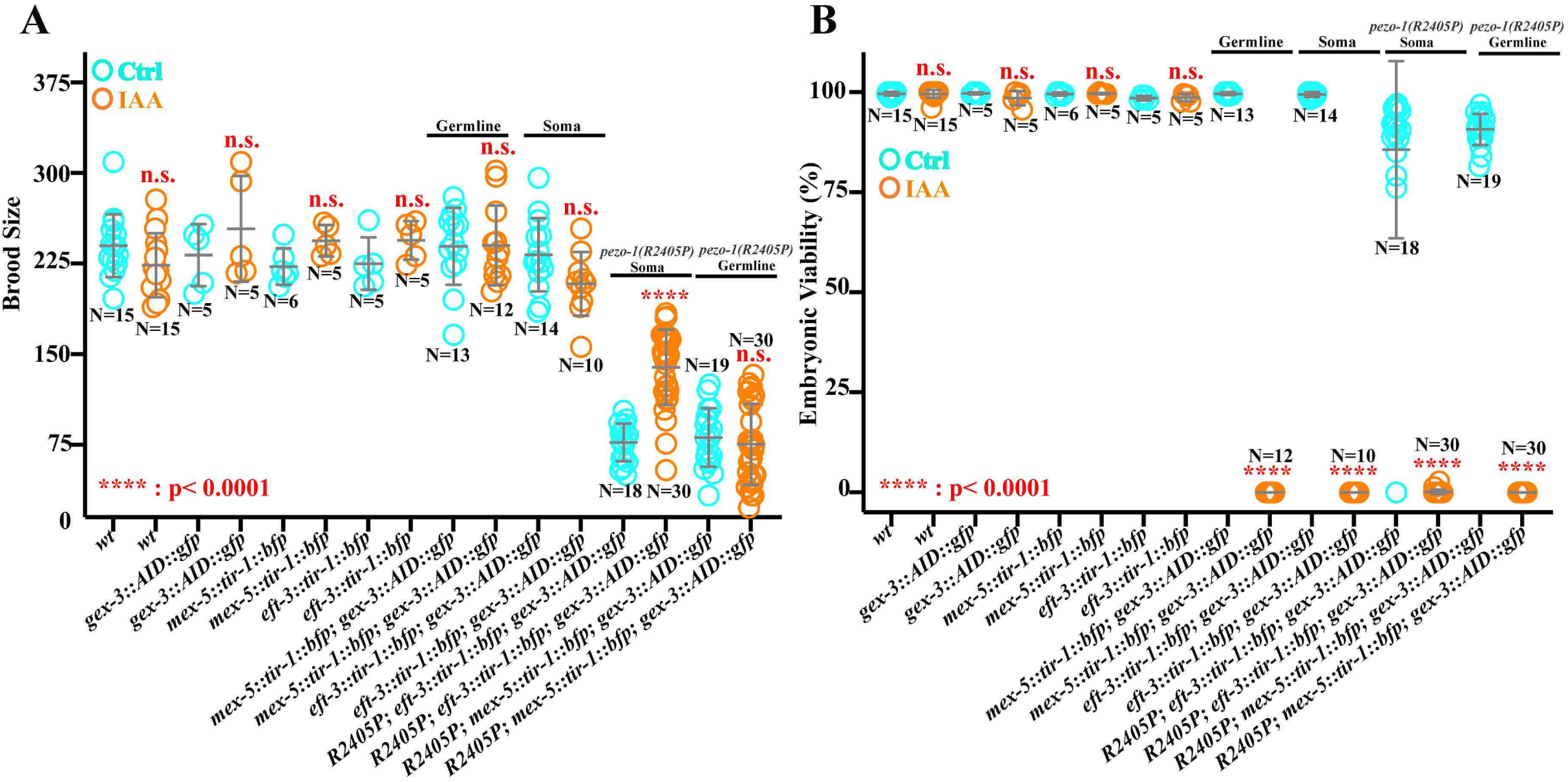
Somatic tissue-specific degradation of GEX-3 suppresses the small brood size in the *pezo-1(R2405P)* mutant. A degron and GFP cassette was inserted at the 5′ end of the *gex-3* coding sequence using CRISPR/Cas9-mediated editing. The transgenic TIR-1::BFP::AID was driven by the *eft-3* promoter to express in most or all somatic tissues, including the spermatheca and the somatic sheath cells. TIR-1::BFP::AID was driven by the germline-specific promoter *mex-5,* which drives expression in the germline and oocytes. (A) Brood size was partially restored in the *pezo-1(R2405P)* degron driven by the *eft-3* promoter when animals were treated with 2 mM auxin. (B) Embryonic viabilities were reduced to nearly zero in all degron strains when treated with 2 mM auxin. N values indicated the number of the tested animals in (A-B). P-values: ****, p<0.0001 (t-test). The blue circles represent those animals treated with the ethanol only control and the orange circles represent those treated with auxin.

### Actin orientation and bundling are affected in *pezo-1* mutants

The WAVE regulatory complex is essential for the organization of the actin cytoskeleton and its dynamics. Therefore, we predicted that *gex-3* may coordinate with *pezo-1* actin organization and orientation in the spermatheca. Previous studies indicated that spermathecal contractility was tightly associated with proper actin organization ^13–15^. To further investigate the functional contribution of *pezo-1* and *gex-3* to spermathecal contractility and actin organization, we used a spermatheca-specific GFP marker GFP::ACT-1 for labeling actin. In mature wild-type animals, the entire actin network was tightly compacted in the contracting spermathecal cells (Fig. 8A-A’). In the dilated spermatheca occupied by an ovulating oocyte, each spermathecal cell contains prominent parallel actin bundles at the cellular cortex which run along the basal cell edges, termed peripheral actin bundles (Fig. 8B-B’). To test whether actin organization was affected by *pezo-1* and *gex-3*, we crossed the actin reporter strain into the following four strains: a) *pezo-1(R2405P)*, b) *pezo-1* full deletion mutant *pezo-1*Δ, c) *gex-3(L353F)*, and d) the *pezo-1(R2405P) gex-3(L353F)* double mutant. We found a variety of defects in actin bundle distribution and orientation in the spermathecal cells of the *pezo-1(R2405P)* and *pezo-1*Δ mutants (Fig. 8C-E’, H-J). In these cells with actin bundle orientation defects the actin bundles within the cells were aligned, but ran perpendicular to the long cell axis, termed perpendicular actin (Fig. 8C-D’, I). We occasionally found that the actin bundling was missing in the spermathecal cells when PEZO-1 was disrupted, which we defined as missing actin (Fig 8C-C’, and J). We also observed aligned actin that was densely associated with thicker and brighter actin-bunches in *pezo-1(R2405P)* and *pezo-1*Δ, which were defined as bunched actin (Fig. 8D-E’, H) ^27^. The *gex-3(L353F)* mutant only caused mild actin defects, including bunching and perpendicular defects (Fig. 8F-F’, H). The *pezo-1(R2405P) gex-3(L353F)* double mutant significantly rescued bunching actin defects but did not affect perpendicular and missing actin (Fig. 8G-G’, I-J). Collectively, these results indicate that *pezo-1* is critical to spermathecal contractility, likely through influencing actin cytoskeletal organization and orientation. *gex-3(L353F)* partially suppressed the actin defects which may contribute to the alleviated ovulation defects and brood size (Fig. 9).

**Figure 8.**
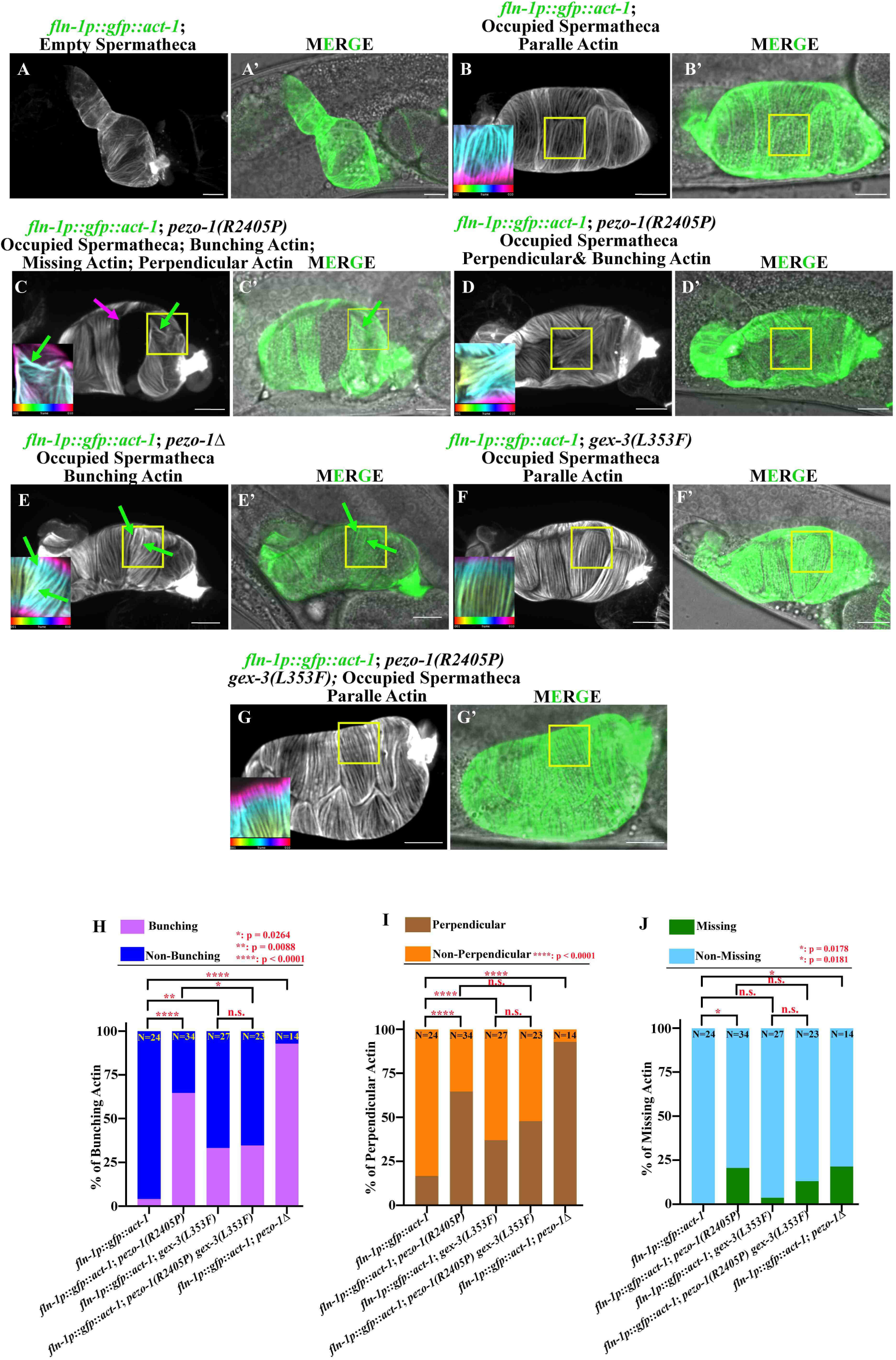
Actin organization and orientation were disrupted in *pezo-1* mutants. (A-A’) Representative images of the contracted spermatheca labelled by the actin marker GFP::ACT-1. (B-B’) Representative images of WT spermathecal cells with parallel actin bundles. The magnified insert from the area in B and B’ are indicated by a yellow box. (C-D’) Representative images of defective actin bundles including perpendicular actin, missing actin, and bunching actin in the *pezo-1(R2405P)* mutant. Bunching actin is indicated by green arrows. Missing actin is indicated by magenta arrows. (E-E’) Representative images of defective actin bundles in *pezo-1*Δ mutant. Bunching actin are indicated by green arrows. (F-F’) Actin organization in *gex-3(L353F)* animals. (G-G’) Double mutant *pezo-1(R2405P) gex-3(L353F)* suppressed the actin defects when compared to the *pezo-1(R2405P)* single mutant. (H-J) Quantification of actin defects in each strain. Insert is color-coded according to z-depth (color scale bar shown on the bottom of panel B, C, D, E, F, and G) to indicate the bundle organization and orientation. N values indicated the number of the tested occupied spermatheca in (H-J). Scale bar was labeled in each panel. P-values: *, p=0.0264 (H); * p= 0.0178 (J); * p= 0.0181 (J); ** p=0.0088 (H); **** p < 0.0001 (H-I) (Chi-squared test).

**Figure 9.**
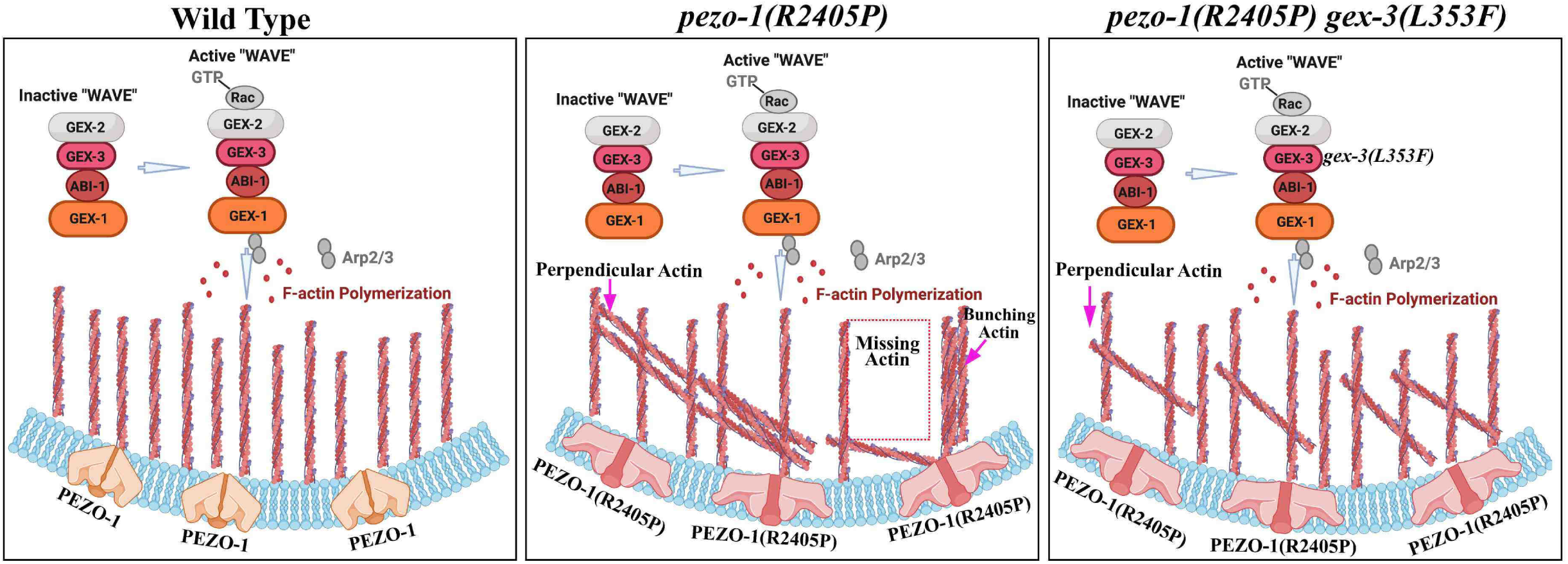
Working model for genetic interaction between *pezo-1(R2405P)* and *gex-3(L353F)* The proposed model suggests how *gex-3(L353F)* or partially depletion of *gex-3* by RNAi might suppress the reproductive deficiency in the *pezo-1(R2405P)* mutant. At the cellular level, the *pezo-1(R2405P)* allele disrupted the actin organization and orientation, such as perpendicular actin, bunching acting, and missing actin in the spermatheca cells. *gex-3(L353F)* allele partially alleviated the actin defects in the *pezo-1(R2405P)* mutant, which may explain its suppression of the low ovulation rate and number of crushed oocytes during ovulation, all leading to suppression of the reduced brood size caused by the *pezo-1(R2405P)* mutant. The proteins labeled by grey color were not explored in this paper.

## Discussion

The PIEZO proteins are associated with at least 26 human disorders and diseases ^3^. More than 100 variants of PIEZO have been identified to cause physiological disorders. Therefore, it is critical to understand the molecular mechanism whereby dysfunctional PIEZO alters physiological processes, as well as identify molecular and genetic determinants that may affect PIEZO activity. Complete knockout of *Piezo1* and *Piezo2* in mice causes embryonic lethality and fetal cardiac defects ^28, 29^. The PIEZO mutants in other systems, such as *Drosophila* or zebrafish, only lead to mild phenotypes ^30–32^, which limited the possibility to perform forward genetic screens to identify genetic determinants of PIEZO in those systems *in vivo*. Our previous study demonstrated that PEZO-1 channel influenced a series of reproductive processes, including ovulation, the expulsion of the fertilized oocyte into the uterus, and sperm navigation ^18^. Dysfunction of PEZO-1 causes a severe reduction in the ovulation rate, defective sperm navigation behavior, and small brood size. The severe reduction in brood size in the *pezo-1* mutants provided an easy and reproducible readout for a forward genetic screen.

*Caenorhabditis elegans* is a powerful model in which gene editing and behavior are becoming an attractive system for precision modeling of human genetic diseases. In this study, we tested five PIEZO pathogenic variants in the *C. elegans pezo-1* gene, all of which displayed similar ovulation phenotypes as our *pezo-1* deletion mutants, suggesting that these alleles compromise PEZO-1 function and/or channel activity. All five variants were localized to the predicted pore of the PEZO-1 channel (Fig. 1A). The conserved human variants cause diseases in various organs or tissues, such as DHS in red blood cells or DA5, and Gordon Syndrome in joint and distal extremities ^3^ (Table 1). The phenotypes observed in *C. elegans* do not reflect these PIEZO-derived disease symptoms, yet our disease modeling pipeline demonstrated the usefulness of the *C. elegans* reproductive tract for investigating the physiological contribution and molecular mechanisms of these PIEZO-based diseases mutations.

The severe reduction in brood size of the pathogenic mutants allowed us to screen putative genetic determinants for PIEZO suppressors *in vivo*. Our chemical mutagen-mediated forward genetic screening combined with MIP-MAP genomic mapping, facilitated the discovery of the suppressor alleles. In this study, we successfully identified the cytoskeletal regulator WAVE/SCAR complex subunit GEX-3, which suppressed the defective phenotypes caused by the gain-of-function *pezo-1(R2405P)* allele. To our knowledge, this is the first genetic suppressor of PIEZO that has being identified through a forward genetic screening approach.

Actin-binding and regulatory proteins are crucial for proper spermatheca contractility and actin organization, which are necessary for achieving proper ovulation ^27^. Our actin imaging data revealed prominent parallel actin bundles, referred to peripheral actin bundles, at the basal cell surface (Fig. 9). In contrast, *pezo-1* mutants disrupted and distorted the actin organization and orientation in the spermatheca during ovulation (Fig. 9), likely contributing to the observed contractility and ovulation defects such as crushed oocytes. Previous studies have indicated that PIEZO channel function relies on the effective communication and physical interactions between the channels and cytoskeletal components, including actin filaments and focal adhesions with the extracellular matrix ^33–37^.

GEX-3 is a component of the WAVE complex, which controls actin cytoskeletal organization and dynamics by triggering the activity of the actin polymerization regulator Arp2/3 complex ^38^. Loss of *gex-3* resulted in a disrupted actin cytoskeleton during embryogenesis and axon migration in *C. elegans* ^39^. Based on our proposed model (Fig. 9), the GEX-3 suppressor allele could affect actin polymerization and actin organization in a way that partially alleviates the actin defects caused by the PEZO-1 gain-of-function mutation, thereby mitigating mechanotransduction deficiencies and improving spermathecal contractility during ovulation. These findings are consistent with recent studies showing that PIEZO activity could enhance actin polarization by physically tethering to the cadherin-β-catenin mechanotransduction complex ^37, 40^ (Fig. 9). Overall, the link between PEZO-1 and the actin cytoskeleton is likely part of a proposed feedback mechanism ^36^. In this mechanism, the activation of PEZO-1 influences the dynamics and formation of cytoskeletal components, while the cytoskeleton affects PEZO-1 activation during mechanical activities like ovulation.

In addition, the suppressor *gex-3* that we identified is likely to be *pezo-1(R2405)* allele specific, as it did not suppress the other four disease alleles generated in this study. We also could not rule out the possibility of an additional suppressor allele within the suppressor background, as we observed that the CRISPR-edited *gex-3* suppressor strain had weaker effects compared to the original suppressor strain. Using this genetic screening pipeline will allow us to identify more suppressors in other pathogenic mutants, as well as uncover the molecular mechanism and functional contribution of each pathogenic allele to PEZO-1 activity. These screens may aid in defining the cellular mechanism that modulate PIEZO channel function and pave the pathway for further therapeutic approaches. We also found that sperm attraction defects were variable or not affected in some *pezo-1* missense mutations, which can allow us to separate the functional studies of sperm attraction signaling pathways from the mechanotransductive processes of ovulation.

Lastly, the human ortholog of *gex-3*, *NCKP1*, plays a critical role in neurodevelopment. Disruptive variants of *NCKP1* have been associated with neurodevelopmental disorders, including Coffin-Siris syndrome 1 (CSS1) and autism spectrum disorder (ASD) ^41, 42^. The underlying causes of the *NCKP1*-associated diseases are likely due to altered actin dynamics, which interfere with neuronal migration during cortical development ^41^. Patients with Coffin Siris syndrome 1 variants exhibit symptoms such as aplasia or hypoplasia of the distal phalanges and abnormal facial features, which are similar to those observed in DA5 patients ^43^. Therefore, by utilizing genetic approaches, we can not only identify suppressors of disease relevant *pezo-1* mutants, but also uncover other molecules that contribute to these symptoms in humans. This helps to establish a molecular connection between disease-causing genes and provides valuable insights for future therapeutic advancements.

## Materials and Methods

### *C. elegans* strains used in this study

*C. elegans* strains were maintained with Golden lab protocols ^44^. Strain information is listed in Table 2.

**Table 2.**
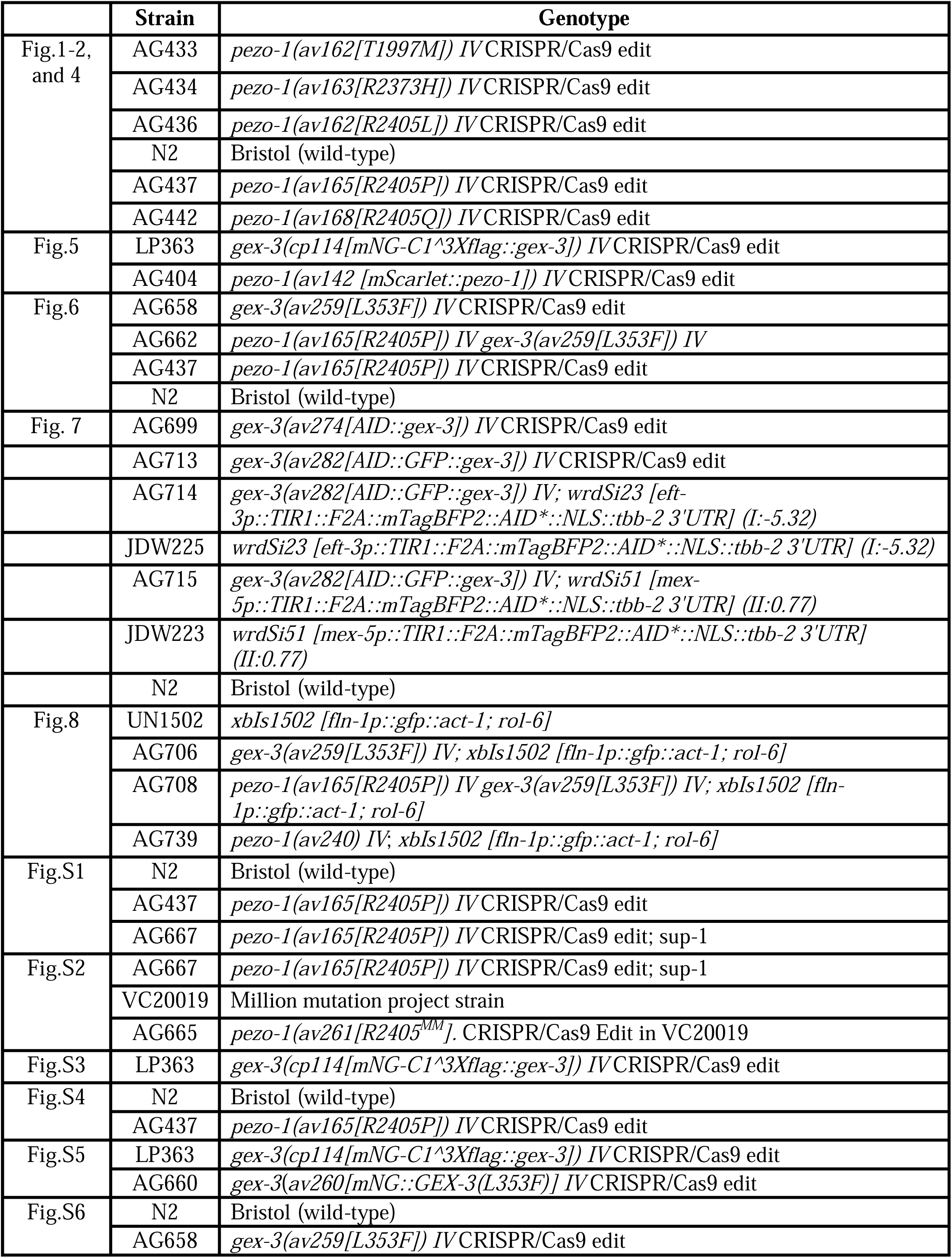
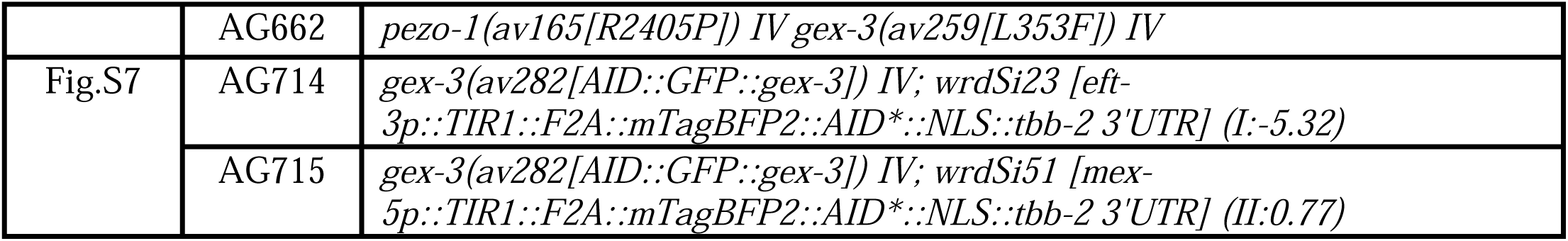
*C. elegans* strains list in the study.

|  | Strain | Genotype |
| --- | --- | --- |
| Fig.1-2,<br>and 4 | AG433 | *pezo-1(av162[T1997M]) IV* CRISPR/Cas9 edit |
|  | AG434 | *pezo-1(av163[R2373H]) IV* CRISPR/Cas9 edit |
|  | AG436 | *pezo-1(av162[R2405L]) IV* CRISPR/Cas9 edit |
|  | N2 | Bristol (wild-type) |
|  | AG437 | *pezo-1(av165[R2405P]) IV* CRISPR/Cas9 edit |
|  | AG442 | *pezo-1(av168[R2405Q]) IV* CRISPR/Cas9 edit |
| Fig.5 | LP363 | *gex-3(cp114[mNG-C1<sup>3</sup>Xflag::gex-3]) IV* CRISPR/Cas9 edit |
|  | AG404 | *pezo-1(av142 [mScarlet::pezo-1]) IV* CRISPR/Cas9 edit |
| Fig.6 | AG658 | *gex-3(av259[L353F]) IV* CRISPR/Cas9 edit |
|  | AG662 | *pezo-1(av165[R2405P]) IV gex-3(av259[L353F]) IV* |
|  | AG437 | *pezo-1(av165[R2405P]) IV* CRISPR/Cas9 edit |
|  | N2 | Bristol (wild-type) |
| Fig. 7 | AG699 | *gex-3(av274[AID::gex-3]) IV* CRISPR/Cas9 edit |
|  | AG713 | *gex-3(av282[AID::GFP::gex-3]) IV* CRISPR/Cas9 edit |
|  | AG714 | *gex-3(av282[AID::GFP::gex-3]) IV; wrdSi23 [eft-3p::TIR1::F2A::mTagBFP2::AID\*::NLS::tbb-2 3'UTR] (I:-5.32)* |
|  | JDW225 | *wrdSi23 [eft-3p::TIR1::F2A::mTagBFP2::AID\*::NLS::tbb-2 3'UTR] (I:-5.32)* |
|  | AG715 | *gex-3(av282[AID::GFP::gex-3]) IV; wrdSi51 [mex-5p::TIR1::F2A::mTagBFP2::AID\*::NLS::tbb-2 3'UTR] (II:0.77)* |
|  | JDW223 | *wrdSi51 [mex-5p::TIR1::F2A::mTagBFP2::AID\*::NLS::tbb-2 3'UTR] (II:0.77)* |
|  | N2 | Bristol (wild-type) |
| Fig.8 | UN1502 | *xbIs1502 [fln-1p::gfp::act-1; rol-6]* |
|  | AG706 | *gex-3(av259[L353F]) IV; xbIs1502 [fln-1p::gfp::act-1; rol-6]* |
|  | AG708 | *pezo-1(av165[R2405P]) IV gex-3(av259[L353F]) IV; xbIs1502 [fln-1p::gfp::act-1; rol-6]* |
|  | AG739 | *pezo-1(av240) IV; xbIs1502 [fln-1p::gfp::act-1; rol-6]* |
| Fig.S1 | N2 | Bristol (wild-type) |
|  | AG437 | *pezo-1(av165[R2405P]) IV* CRISPR/Cas9 edit |
|  | AG667 | *pezo-1(av165[R2405P]) IV* CRISPR/Cas9 edit; sup-1 |
| Fig.S2 | AG667 | *pezo-1(av165[R2405P]) IV* CRISPR/Cas9 edit; sup-1 |
|  | VC20019 | Million mutation project strain |
|  | AG665 | *pezo-1(av261[R2405<sup>MM</sup>]. CRISPR/Cas9 Edit in VC20019* |
| Fig.S3 | LP363 | *gex-3(cp114[mNG-C1<sup>3</sup>Xflag::gex-3]) IV* CRISPR/Cas9 edit |
| Fig.S4 | N2 | Bristol (wild-type) |
|  | AG437 | *pezo-1(av165[R2405P]) IV* CRISPR/Cas9 edit |
| Fig.S5 | LP363 | *gex-3(cp114[mNG-C1<sup>3</sup>Xflag::gex-3]) IV* CRISPR/Cas9 edit |
|  | AG660 | *gex-3(av260[mNG::GEX-3(L353F)]) IV* CRISPR/Cas9 edit |
| Fig.S6 | N2 | Bristol (wild-type) |
|  | AG658 | *gex-3(av259[L353F]) IV* CRISPR/Cas9 edit |
|  | AG662 | *pezo-1(av165[R2405P]) IV gex-3(av259[L353F]) IV* |
| Fig.S7 | AG714 | *gex-3(av282[AID::GFP::gex-3]) IV; wrdSi23 [eft-3p::TIR1::F2A::mTagBFP2::AID\*::NLS::tbb-2 3'UTR] (I:-5.32)* |
|  | AG715 | *gex-3(av282[AID::GFP::gex-3]) IV; wrdSi51 [mex-5p::TIR1::F2A::mTagBFP2::AID\*::NLS::tbb-2 3'UTR] (II:0.77)* |

### EMS Suppressor Screen

AG437 *pezo-1(R2405P)* early L4 hermaphrodites were washed three times in M9 and soaked in 48 mM Ethyl methane sulfonate (EMS) solution for four hours at room temperature. The EMS treated animals were washed three times in M9 and were transferred to a fresh 100 mm MYOB plate with OP50 on one side. The animals were allowed to recover up to 4 hours before being picked to individual 100 mm MYOB plates with fresh OP50. Only the recovered animals that were able to crawl across the plates to the OP50 food were transferred to the fresh plates. A total of 70 MYOB plates with 25-30 mid-L4 (P0s) on each were incubated at 20 °C. Gravid F1 adult were bleached and F2 embryos were collected after hypochlorite treatment. The F2s embryos were shaken in a glass flask with M9 buffer overnight, and hatched larvae were grown on *sca-1* RNAi plates at 25 °C for a week. Their progeny was screened for viable larvae. ∼150k mutagenized haploid genomes were scored in this fashion.

### RNAi treatment

The RNAi feeding constructs were chosen from the Ahringer and Vidal libraries ^45, 46^. RNAi bacteria were grown until log phase was reached and spread on MYOB plates containing 1mM IPTG and 25 μg/ml carbenicillin and incubated 12-14 hours. The seeded RNAi plates were stored at 4 °C up to one week. To deplete the target genes *gex-3*, *gex-1*, and *abi-1*, mid-L4 hermaphrodites were picked onto plates with the IPTG-induced bacteria. Animals were grown on RNAi plates at 20°C for 36-60 hours for brood size and other assays.

### Brood size determinations and embryonic viability assays

Single mid-L4 hermaphrodites were picked onto 35 mm MYOB plates seeded with 5-10 μl of fresh OP50 bacteria and allowed to lay eggs for 36 hours (plate one contains the brood size from 0-36 hours post mid-L4). The hermaphrodites were transferred to a newly seeded 35 mm MYOB plate to lay eggs for another 24 hours and were flamed from the plate (the brood size on this plate defined as the brood size from 36-60 hours post mid-L4). Twenty-four hours after removing the hermaphrodites, the viable larvae were counted for the embryonic viability. Brood sizes were determined at 36 hours and 60 hours. Percentage of embryonic viability= (the number of hatched larva / the total brood size) *100%.

### Live imaging to determine ovulation rates

For imaging ovulation, animals were immobilized on 7% agar pads with anesthetic (0.1% tricaine and 0.01% tetramisole in M9 buffer). DIC image acquisition was captured by a Nikon 60 X oil objective with 2-3 μm z-step size; 15-25 z planes were captured. Time interval for ovulation imaging is every 60-90 seconds, and duration of imaging is 60-90 minutes. Ovulation rate= (number of successfully ovulated oocytes) / total image duration. Actin imaging were captured by a Nikon 60X water objective with 0.5 μm z-step size; 15-20 z planes were captured.

### CRISPR design

All CRISPR/Cas9 editing was generated into Bristol N2 strain as the wild type unless was described separately. The crRNAs were synthesized from Horizon discovery, along with tracrRNA. Repair template and oligos were purchased from Integrated DNA Technologies (IDT). The CRISPR design followed the standard protocols ^47^. Approximately 20-30 young gravid animals were injected with the CRISPR/Cas9 injection mix. Detailed sequence information of CRISPR design were listed below: (Capital letters represent the ORF or exon sequence, lowercase letters indicate the sequence from intron). Detailed information is listed in Table 3.

**Table 3.**
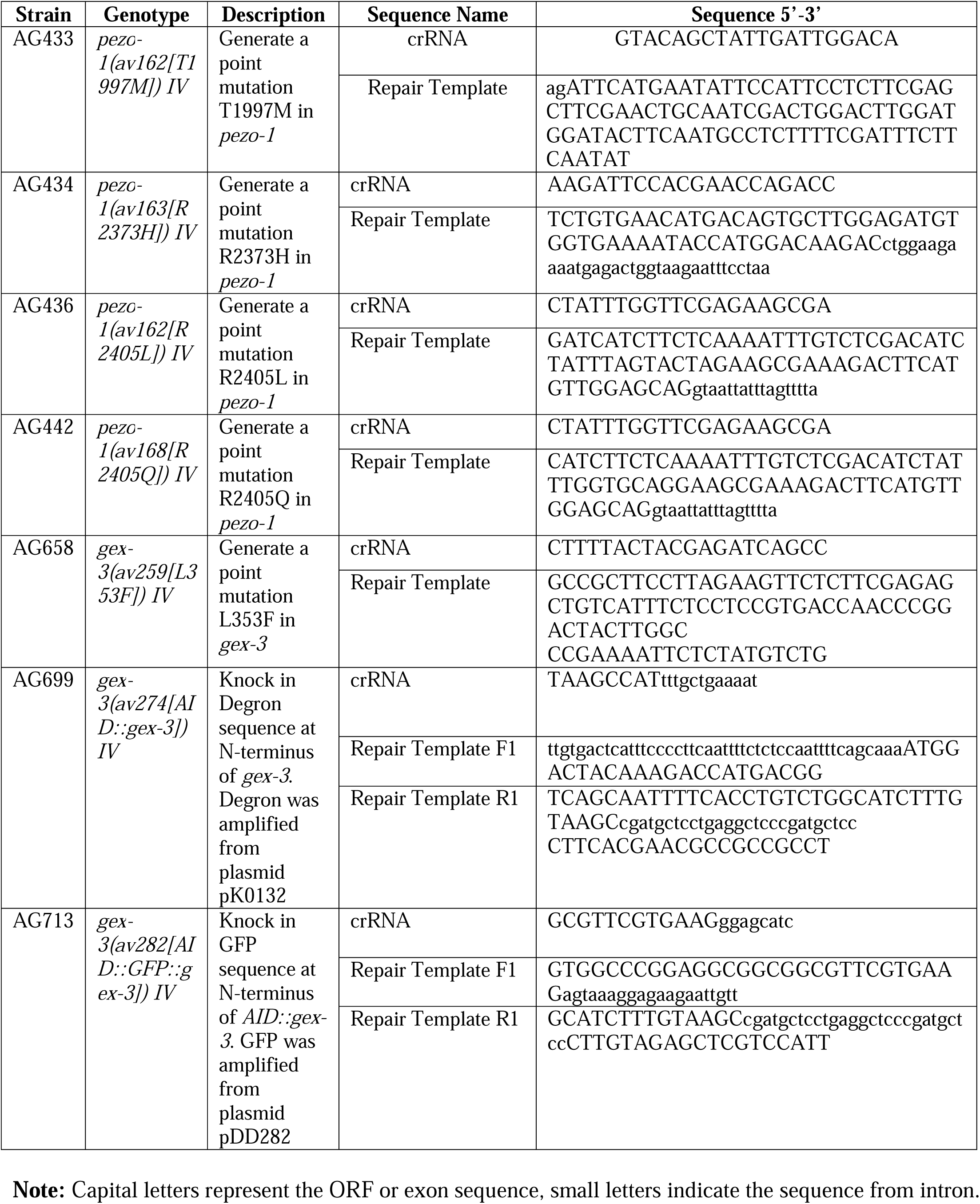
List of the sequence for the CRISPR design.

### Sperm distribution assay and mating assay

MitoTracker Red CMXRos (MT) (Invitrogen # M7512) was used to label male sperm following the protocol adapted from previous studies ^17, 48^. Wild type males were incubated in the MT buffer for 2 hours in the dark. The stained males were covered by foil to prevent light exposure overnight. About 30 males were placed with 10 anesthetized hermaphrodites (0.1% tricaine and 0.01% tetramisole in M9 buffer) on MYOB plates seeded with a 10 μl OP50 bacteria. After 20-30 minutes of mating, hermaphrodites were then isolated and allowed to rest on food for at least one hour. The mated hermaphrodites were then mounted for microscopy on 7% agarose pads with the anesthetic. Image acquisition was captured by a Nikon 60 × 1.2 NA water objective with 1 um z-step size. Sperm distributions were quantified by Image J-cell counter.

### Auxin-inducible treatment in the degron strains

Auxin indole-3-acetic acid (IAA) was purchased from Alfa Aesar (#A10556). A 400 mM stock solution of IAA was made in ethanol and was added to MYOB medium to a final concentration of 1 or 2 mM auxin. To efficiently degrade the GEX-3 protein, mid-L4 hermaphrodites were picked onto auxin plates. Animals were grown on the plates at 20°C for 24 hours for the degradation efficiency test, and 60 hours for brood size assay.

### Microscopy

All imaging were performed on a spinning disk confocal system that uses a Nikon 60 × 1.2 NA water or oil objectives, a Photometrics Prime 95B EMCCD camera, and a Yokogawa CSU-X1 confocal scanner unit. Nikon’s NIS imaging software were applied to capture the images. The image data were processed using ImageJ/FIJI Bio-formats plugin (National Institutes of Health)^49, 50^.

### Expression of *pezo-1* in Sf9 insect cells

To express PEZO-1 in Sf9 cells (a clonal isolate of *Spodoptera frugiperda* Sf21 cells), we generated recombinant baculoviruses, according to the manufacturer’s instructions (Bac-to-Bac expression system; Invitrogen). To generate this baculovirus, we used a pFastBac construct (Epoch Life Science) containing an 8× histidine–maltose binding protein tag and a synthesized *pezo-1* isoform G nucleotide sequence (one of the longest isoforms according to RNA sequencing; wormbase.org release WS280). For expression of PEZO-1 R2405P, the construct contained an 8× histidine–maltose binding protein tag and a synthesized *pezo-1* isoform G with the R2405P point mutation. We infected Sf9 cells with either wild-type or mutant *pezo*-*1* baculovirus for 48 h as previously established (Millet et al, 2022). Infected cells were plated on glass coverslips coated with a peanut lectin solution (1 mg/ml; Sigma-Aldrich) for patch-clamp experiments.

### Electrophysiology and mechanical stimulation

Control and *pezo-1* (wild-type or R2405P constructs) infected Sf9 insect cells were recorded in the whole-cell patch-clamp configuration, as previously described (Millet et al, 2022). The bath solution contained 140 mM NaCl, 6 mM KCl, 2 mM CaCl2, 1 mM MgCl2, 10 mM glucose, and 10 mM HEPES, pH 7.4. The pipette solution contained 140 mM CsCl, 5 mM EGTA, 1 mM CaCl2, 1 mM MgCl2, and 10 mM HEPES, pH 7.2. Sf9 cells were mechanically stimulated with a heat-polished blunt glass pipette (3–4 µm) driven by a piezo servo controller (E625; Physik Instrumente). The blunt pipette was mounted on a micromanipulator at an ∼45° angle and positioned 3–4 µm above the cells without indenting them. Displacement measurements were obtained with a square-pulse protocol consisting of 1-µm incremental indentation steps, each lasting 200 ms, with a 2-ms ramp in 10-s intervals. Recordings with leak currents >200 pA and with access resistance >10 MΩ, as well as cells with giga-seals that did not withstand at least five consecutive steps of mechanical stimulation, were excluded from analyses. Pipettes were made from borosilicate glass (Sutter Instruments) and were fire-polished before use until a resistance between 3 and 4 MΩ was reached. Currents were recorded at a constant voltage (−60 mV), sampled at 20 kHz, and low-pass filtered at 2 kHz using a MultiClamp 700B amplifier and Clampex (Molecular Devices). Leak currents before mechanical stimulation were subtracted offline from the current traces. The time constant of inactivation τ was obtained by fitting a single exponential function, Eq. 1, between the peak value of the current and the end of the stimulus:

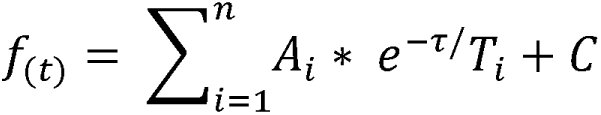

where A = amplitude, τ = time constant, and the constant y-offset C for each component i.

### MIP-MAP and Data analysis

*pezo-1(R2405P^MM^)* males were mated with the homozygous hermaphrodites of each suppressor line (Fig. S2A). We carefully pooled suppressed F2 progeny from the cross and allowed the F2 population to expand for ten generations for MIP-MAP analysis. Ten generations of self recombination are sufficient to distribute the MIP-MAP single nucleotide polymorphism (SNPs) (also refer to molecular probes) into the suppressor line background and provide a high molecular resolution to identify the mutated regions. Candidate mutations (defined as novel, homozygous, and nonsynonymous) were identified by whole-genome sequencing as described previously ^51^.

Briefly, sequencing libraries were constructed by Invitrogen Pure Link Genomic DNA Mini Kit (Ref # K1820-01) with genomic DNA from homozygous suppressor-bearing strains. The libraries were pooled and sequenced on a HiSeq 3000 instrument (Illumina, San Diego, CA) to at least 20-fold coverage. Variants were identified with a pipeline of BBMap [56], SAMtools ^52^, FreeBayes [57], and ANNOVAR [58]. Mapping loci for suppressors were identified using molecular inversion probes (MIPs) to single-nucleotide polymorphisms (SNPs) as described previously ^20^. Briefly, suppressor-bearing strains were mated to SNP mapping strain VC20019 ^53^ which had been engineered via CRISPR to contain the *pezo-1(R2405P)* mutation. F1 cross progeny were allowed to self-fertilize, and a minimum of 50 homozygous F2 progeny were pooled for construction of MIP libraries. SNP allele frequencies were determined using a custom script and plotted with R [59] to delimit the mapping interval.

### Data and statistical analyses

The electrophysiological data and statistical analyses were performed using GraphPad Instat 3, and OriginPro 2018 software. Statistical methods and sample numbers are detailed in the corresponding figure legends. No technical replicates were included in the analyses. Statistical significance for other assays was determined by p value from an unpaired 2 tailed t-test, one-way ANOVA, or Chi-squared-test. P-values: ns = not significant; * = <0.05; ** = <0.01, *** = <0.001; **** = <0.0001. Both the Shapiro-Wilk and Kolmogorov-Smirnov Normality test indicated that all data follow normal distributions.

## Acknowledgments

We thank the *Caenorhabditis* Genetics Center, which is funded by National Institutes of Health Office of Research Infrastructure Programs (P40OD010440), for providing strains for this study. We thank Dr. Erin Cram’s generosity for providing the actin reporter strains. We are grateful to the members of the Golden laboratory for productive discussions and preparing reagents. We especially thank Dr. Erin Cram, Dr. Orna Cohen-Fix, Dr. Kevin O’ Connell, and Dr. Katherine McJunkin for critical inputs on the project and feedback on the manuscript. We thank all members of the Baltimore Worm Club for providing feedback and suggestions to our investigations. In Memoriam of Dr. Andy Golden, who provided tremendous both scientific and financial support to this study. He will be sorely missed.

---

**Supplemental Figure 1 Forward genetic screen for suppressors of the small brood size of a DA5 patient-specific allele *pezo-1(R2405P)*** (A) Strategy to identify genetic modifiers that restore the small brood size of *pezo-1(R2405P)* mutants. The synchronized L4 larvae of the *pezo-1(R2405P)* mutation (P0) were collected and treated with EMS. The healthy EMS-treated late L4 larvae were picked to a single plate to propagate F1 populations at 20°C. The gravid F1 adults were bleached, and the viable F2 larvae were transferred to *sca-1* RNAi plates at 25°C. These F2 animals were allowed to grow and were screened for fertility and restored brood sizes. (B) Brood size of wild type, *pezo-1(R2405P)*, and the suppressor line at different conditions over 60 hours post mid-L4. N values indicated the number of the tested animals in (B). P-values: ****, p<0.0001 (t-test). The graphic was generated with BioRender.com.

**Supplemental Figure 2 Mapping of *pezo-1(R2405P)* suppressor via VC20019 and MIP-MAP sequencing.** (A) Diagram of the MIP-MAP workflow to map the single nucleotide variants (SNVs) in the suppressor line. (B-F) The read frequency of the VC20019-specific SNVs across the genome of the pooled F2 progeny from *pezo-1(R2405P)*; *sup1* and *pezo-1(R2405P)*^MM^ cross. (B) A mutation-associated interval was identified on Chromosome IV. The strategy graphic was generated with BioRender. com.

**Supplemental Figure 3 *gex-3* RNAi is specific and efficient to reduce the expression of *gex-3* in the germline and somatic tissue** (A-A’’) mNG::GEX-3 (green in A’’) was strongly expressed on germline and oocyte membranes, and in the spermatheca (red arrowheads in A-A’’). (B-B’’) Depletion of *gex-3* by RNAi significantly reduced the expression of mNG::GEX-3 on the germline and oocyte membranes and in the spermatheca. DIC images are shown in panels A’ and B’. (C) Quantification of fluorescent intensity of both germline and spermathecal mNG::GEX-3 with ctrl and *gex-3* RNAi. N values indicated the number of the quantified animals in (C). Scale bars are indicated in each panel. P-values: ****, p<0.0001 (t-test).

**Supplemental Figure 4 Other WAVE complex subunits did not suppress the small brood size in *pezo-1(R2405P)* mutants.** (A-B) 0-60 hours brood size of wild type and *pezo-1(R2405P)* animals after feeding control, *abi-1* (A), and *gex-1* (B) RNAi. N values indicated the number of the tested animals in (A-B)

**Supplemental Figure 5 mNG::GEX-3(L353F) did not disrupt the cellular localization of mNG::GEX-3.** mNG::GEX-3(L353F) (B and D) presents an identical expression pattern as mNG::GEX-3(+) in a variety of cell types, including the pharyngeal-intestinal valve (red arrows in A-B), germline cells and spermathecal cells (yellow arrows in C-D). Scale bars are indicated in each panel.

**Supplemental Figure 6 *gex-3(L353F)* did not rescue sperm attraction defects in the *pezo-1(R2405P)* mutant.** (A-A’) Sperm distribution was not affected in the *gex-3(L353F)* mutant. (B-B’) Double mutant *pezo-1(R2405P) gex-3(L353F)* showed sperm attraction defects identical to *pezo-1(R2405P)* (Fig.2 E-E’). (C) Quantification of sperm distribution values of *gex-3(L353F)* and *pezo-1(R2405P) gex-3(L353F)* double mutant. P-values: ****, p<0.0001 (t-test). Yellow asterisks indicate the vulva (A’ and B’). Scale bars are indicated at top right in each panel.

**Supplemental Figure 7 Tissue-specific degradation of GEX-3 displays a reduced mNG::GEX-3 fluorescence in each tissue expressing TIR-1::mTagBFP2::AID.** (A–D′′) AID::mNG::GFP::GEX-3 localized to reproductive tissues, such as the plasma membrane of the germline cells, oocytes, and spermatheca (red arrows, A, A′′, C, C’’). (A–A′′) TIR-1::mTagBFP2::AID driven by the germline-tissue-specific promoter *mex-5* is strongly expressed in the germline and oocyte nuclei (B, B’’) Fluorescent signals of AID::mNG::GEX-3 at the germline and oocyte membrane were significantly reduced when animals were treated with 2 mM auxin. However, the fluorescent signals of AID::mNG::GEX-3 at somatic spermatheca and sheath cells are not affected (B-B’’). The signal of TIR-1::mTagBFP2::AID was diminished when exposed to the auxin, regardless of the promoters were used to drive its expression (A’, B’, C’, and D’). (C-C’’) TIR-1::mTagBFP2::AID driven by the somatic specific promoters *eft-3* is strongly expressed in the nuclei of all or most somatic tissues. (D-D’’) Fluorescent signals of AID::mNG::GEX-3 driven by *eft-3* promoter at spermatheca and somatic sheath cells are significantly reduced when animals were treated with 2 mM auxin. However, fluorescent signals of AID::mNG::GEX-3 driven at germline cells, oocyte and sperm are not affected. (E–F) Quantification of the fluorescent signals of AID::mNG::GEX-3 under the different conditions. N values indicated the number of the quantified regions in (E-F). P-values: ****, p<0.0001 (t-test).

